# A Frontal-Sensory Cortex Circuit Encoding Context-Dependent Stimulus Statistics

**DOI:** 10.64898/2026.05.12.724533

**Authors:** Tal Dalal, Rafi Haddad

## Abstract

Animals continuously estimate the statistics of sensory events in their environment to guide behavior. Yet, how the brain constructs internal models that represent the probability distributions of stimuli through learning, how these models are updated during rapid contextual shifts, and what brain regions integrate contextual and probabilistic information remains unclear. Here, we devised a behavioral task in which mice used rapidly changing contextual information to estimate the probability of upcoming odors. Recordings from the piriform and orbitofrontal cortices during learning revealed two distinct neuronal subpopulations that differentially encoded odor probabilities. One subpopulation exhibited ramping activity that scaled with the estimated probability of the behaviorally relevant odor and sharply dropped at its onset. In contrast, a second subpopulation became active at odor onset and was modulated by prediction. Notably, probability information emerged in the orbitofrontal cortex before the piriform cortex, and silencing the orbitofrontal cortex abolished the propagation of this information to the piriform cortex, impairing probability learning and retrieval but not context identification. These findings reveal a frontal–sensory circuit that facilitates the learning and representation of context-dependent, behaviorally relevant event probabilities, and suggest that learned state transitions, a core feature of a cognitive map, are propagated to the sensory cortex through learning.

## Introduction

To adapt and survive in a rapidly changing environment, animals need to estimate the probability of future sensory events. This requires the formation of internal models that represent context-dependent event probabilities, which can be flexibly retrieved in different contexts. While the neural coding of reward probability has been extensively studied ^1–3^, how context-dependent stimulus probabilities are encoded has received significantly less attention ^4–6^. Specifically, how neural circuits evolve through learning to represent probability density functions of stimuli, how they are retrieved upon rapid contextual changes, and which brain regions encode this contextual and probabilistic information is largely unknown.

Recent theoretical work ^7^, supported by experimental evidence ^4–6,8^, has shown that prior knowledge can be integrated with sensory evidence already in the sensory cortex. Moreover, several studies provided evidence of predictive activity in a sensory cortex before the stimulus occurs ^9–13^, and reported response modulations upon prediction violations ^5,8,14–21^. These activity patterns were shown to be behaviorally relevant ^11,18,21^, and support the predictive processing framework, which postulates that top-down predictions from higher brain regions are fed to regions lower in the hierarchy to generate predictive activity and prediction-error (PE) signals ^22,23^. Despite these insights, it remains unclear how learned context-dependent event likelihoods are incorporated into predictive activity and PE signals.

To address these questions, we devised a probability-based odor-discrimination task (CS+ vs CS- odors), in which the probability of CS odor occurrence depends on the context, which can change on a trial-by-trial basis. This design facilitates simultaneous learning of explicit, reinforced odor-reward associations (i.e., CS+ predicts reward) and inference of latent odor probabilities (i.e., the context-dependent probabilities of CS+ and CS-). Critically, we designed this paradigm such that context is explicitly cued on each trial to drive context-dependent learning of odor statistics, and we required a behavioral marker, such as reaction time or false alarm rate, to evaluate mice’s internal probability estimates across learning stages. To test where and how learned odor probability information is encoded, we recorded the neural activity in two brain regions throughout task learning: the anterior piriform cortex (aPC), a major part of the olfactory cortex, and the orbitofrontal cortex (OFC), which has been shown to mediate outcome-guided behavior and context-dependent reward associations ^24–27^. The aPC, a three-layer paleocortex, integrates information from multiple associative brain regions, including the OFC ^28–30^, making it an attractive region to study context-dependent stimulus probability encodings. However, it is currently unknown what information the OFC projections convey to the aPC.

We identified two distinct neuronal subpopulations that emerged with learning and encoded probability information differentially. One subpopulation exhibited ramping activity that scaled with the CS+ odor probability and sharply decreased its activity at the CS odor onset. By contrast, a second subpopulation became active at the CS odor onset and signaled prediction violation. These activity patterns were identified in both brain regions, but emerged earlier in the OFC. Using bilateral silencing of OFC activity, we demonstrate that information in aPC about odor probability, but not context identity, is lost when the OFC is silenced, indicating its crucial role in propagating probability-contingency information into sensory-evoked activity patterns. Finally, we demonstrate that the OFC is essential for learning and retrieving odor probability contingencies, but not for learning stimulus-reward associations. These findings shed light on a neural circuit that evolves to represent probability estimations during rapid contextual switches and propagate this information from the frontal cortex to sensory regions during learning.

## Results

### Mice infer and utilize latent context-dependent odor probabilities

To study context-dependent probabilistic learning, we devised a probabilistic odor-discrimination task in which head-fixed mice could estimate the probabilities of the CS+ or CS- odor occurrences in a context-dependent manner (Fig. 1a-b, N = 9 mice). The CS+ odor predicted reward, and the CS- predicted no reward. Each trial started with the presentation of one of two context odors, A or B, which determined the probability of occurrence of the subsequent CS+ or CS- odor (Fig. 1b; number of trials per session N = 144 ± 35, mean ± standard deviation). The CS+ odor was presented in 75% of the trials following the A odor (context A) and 25% following odor B (context B), and vice versa for the CS- odor (Fig. 1c). Within the session, the number of trials in each context and the overall number of CS+ and CS- trials were similar and presented in a pseudo-random order. This task design allowed mice to generate context-dependent probabilistic predictions about the identity of the upcoming odor on a trial-by-trial basis. Two of the nine mice learned an additional context, M, which predicted the upcoming occurrence of CS+ and CS- odors with equal probabilities (Extended Data Fig. 9), and the identity of context odor A and B was switched across mice. Mice rapidly learned to discriminate between the CS+ and CS- odors (Fig. 1d and Extended Data Fig. 1a), and there was no difference in the learning rate between the two contexts (Extended Data Fig. 1b). As learning progressed, we observed higher false alarm rates in the A context, due to the higher predictability of the CS+ odor in this context (Extended Data Fig. 1e). Consistently, mice exhibited higher anticipatory licking rates and faster reaction times to the CS+ in the A compared with the B context (Fig. 1e-f and Extended Data Fig. 1f). These behavioral markers indicate that mice inferred the context-dependent task contingencies, learning that the CS+ odor is more likely to occur in context A than in context B. On average, mice exhibited significantly faster reaction times in the A context compared to the B context when a discrimination task performance was higher than 70%. We therefore defined sessions with a performance level below 70% as novice and above it as expert (Fig. 1g; N = 16 novice and 62 expert sessions). Mice reached the expert state on average after 2.39 ± 0.38 sessions (mean ± SEM; Extended Data Fig. 1c). As mice gained task expertise, lick reaction times were delayed in both contexts, consistent with ^18^; however, mice still licked faster in context A than in context B (Fig. 1g-h and Extended Data Fig. 1g), indicating that mice maintained and utilized the predictive information from the trial context even at high task expertise. We verified that task learning did not depend on other sensory cues (Methods), and faster reaction times did not depend on the odor identity used to signal the A context (Extended Data Fig. 1h). Moreover, sniff rates were highly similar in the two contexts (Extended Data Fig. 1i). A similar trend was observed in mice trained on the three contexts, although the reaction time differences were less pronounced due to a smaller number of trials in each context (Extended Data Fig. 9). In summary, this behavioral paradigm enables the explicit learning of odor discrimination along with implicit inference of latent odor statistics, which is manifested in faster lick reaction times and higher false alarm rates across all learning stages. Mice exhibited fast learning of the odor discrimination task, followed by learning the context-dependent odor statistics (Fig. 1g). We next asked how these odor probability contingencies are encoded in the anterior piriform cortex during learning.

**Fig. 1:**
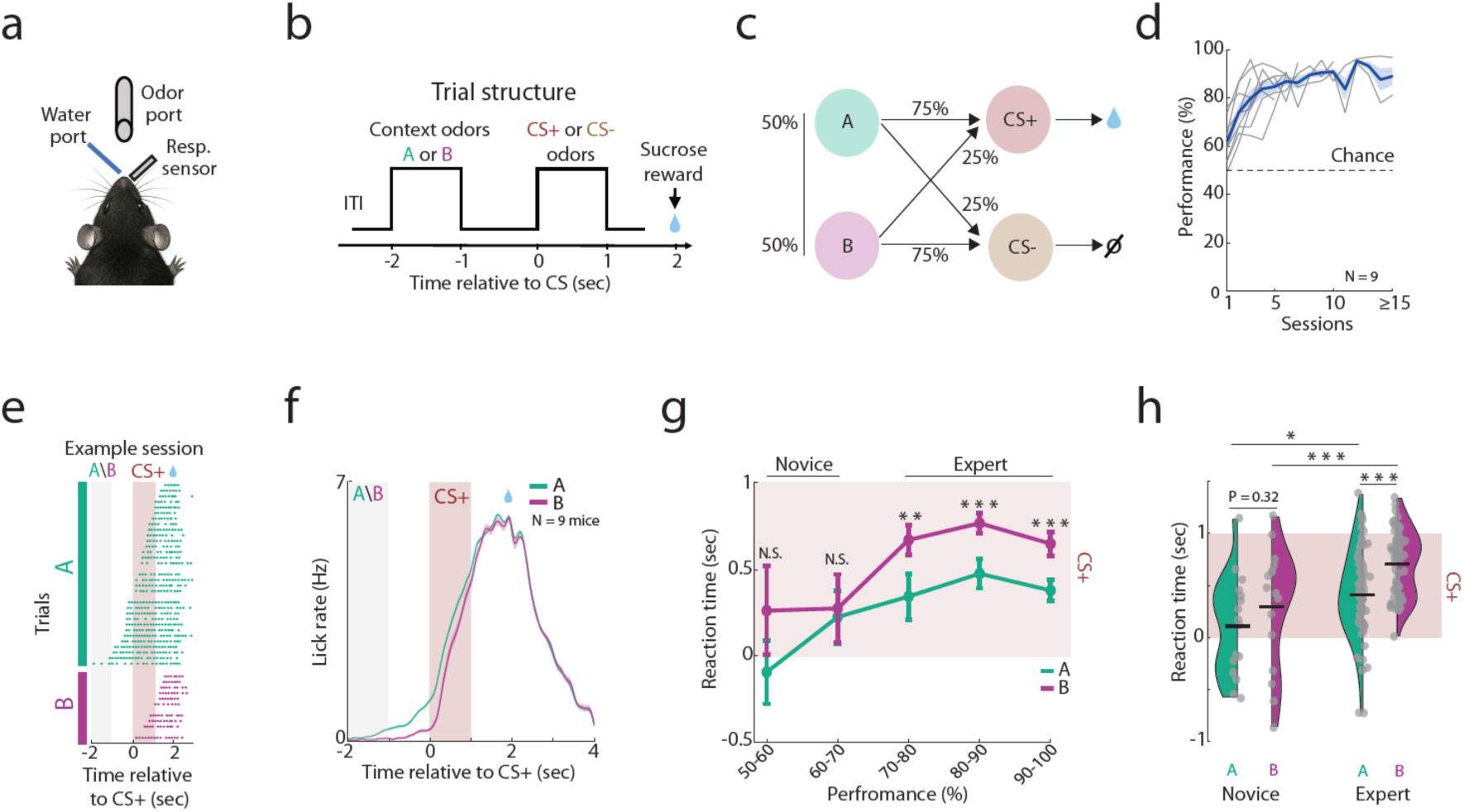
Mice infer and utilize latent context-dependent odor probabilities. a. Schematic illustration of the experimental setup. Head-fixed, water-deprived mice performed a probabilistic odor-discrimination task. Respiratory activity was recorded during task performance. b. Illustration of the behavioral paradigm. Each trial consisted of two consecutive odors. The context odors (A or B) predicted the identity of the subsequent odor (CS+ or CS-) with different probabilities. c. Task probability contingencies. The number of trials in A and B contexts, as well as the number of rewarded (CS+) and unrewarded (CS-) trials, was similar. d. Learning curves of the odor discrimination task across all mice (N = 9, grey curves). The mean ± SEM is shown in blue. e. An example session showing a lick raster plot in the A and B contexts (green and purple, respectively). Reaction times are faster in A compared with the B context (P = 0.0056, two-tailed paired *t*-test). Trials were sorted in each condition by the latency of the first lick. f. Averaged lick PSTHs of hit trials in A and B contexts in expert mice (N = 9 mice; green and purple, respectively; 62 sessions). A higher rate of anticipatory licking is observed before the CS+ odor presentation in the A context compared to the B context. g. Reaction times as a function of the task performance across all sessions. The difference in reaction times between A and B contexts is evident when the discrimination performance reaches 70% (N.S., P > 0.34, ***P < 0.001, **P < 0.01; two-tailed paired *t*-test). The red patch indicates the duration of the CS+ odor. h. Lick reaction time analysis across all mice, showing faster reaction times for the A context in expert but not in novice state (N = 9 mice; novice, N = 16 sessions, P = 0.32; expert, N = 62 sessions, ***P < 1x10^-6^, two-tailed paired *t*-test). Reaction times in both contexts were delayed as mice transitioned to the expert state (*P < 0.05 and ***P < 0.001, two-tailed paired *t*-test comparing reaction times in the A and B contexts for novice and expert states, respectively).

### Ramping activity in aPC encodes the estimated probability of the behaviorally relevant odor

To examine how neural activity in the olfactory cortex encodes learned context-dependent task contingencies, we recorded the spiking activity in the anterior piriform cortex (aPC) throughout the entire learning process using chronically implanted tetrodes (Extended Data Fig. 2a). For reference, we recorded the neural activity in response to passive exposure to the four odors used in our experiment before training began (N = 72 neurons from 6 mice; Extended Data Fig. 2b-g, Methods). Overall, we recorded 401 well-isolated units from four mice during task performance. Two of the four mice were also trained on the M context (P = 0.5 for CS+ and CS-, N = 305 neurons; Extended Data Fig. 9 and Methods). In the subsequent sections, we primarily focus our analyses on A and B contexts for statistical power, and use the neural activity recorded in the M context for additional analyses. Most of the recorded neurons exhibited significant task-modulated activity and mixed selectivity for different parts of the trial (Fig. 2a and Extended Data Fig. 3a-b; N = 316/401 neurons, 79% of all recorded neurons, Methods). We hypothesized that as mice learned to estimate the relative probabilities of the CS odors, changes in neural activity would be prominent before their onset, that is, during the delay period. To investigate this at the population level, we projected the population activity onto the first two principal components, which together accounted for 73% of the variance, as a function of time during the delay period. This process yielded the population trajectory in the neural state space for each context. We then computed the Mahalanobis distance in state space between the population trajectories during the late delay period and baseline, separately for the novice and expert states (Fig. 2b and Extended Data Fig. 3c). We found that population activity strongly diverged before the onset of the CS odor in the A, B, and M contexts. The deviation of these population trajectories from baseline scaled with the relative CS+ odor probability.

These graded dynamics were absent in the novice state, indicating that they emerged following the learning of task contingencies, rather than reflecting residual odor-evoked activity differences between the context odors during the delay period (Fig. 2b). To further investigate these delay period dynamics, we examined whether individual neurons carry context-dependent probability information about the upcoming odors on a trial-by-trial basis. Focusing on the late part of the delay period (-330 ms to 0, as in ^9,10,31^, we identified a subpopulation of neurons that ramped their activity during the delay period up to, or shortly before, the onset of the CS odors (Fig. 2c, ‘ramping neurons’, 23% of task-responsive neurons, 73/316 neurons). Ramping activity either transiently increased before the CS odor onsets (e.g., Fig. 2c, neuron #2) or had a sustained activity throughout the delay period (e.g., Fig. 2c, neuron #4). Ramping activity during the late delay period reliably encoded the trial context (i.e., A or B, using the activity during -330 ms to 0), significantly better than non-ramping neurons (Fig. 2d). Interestingly, the rate of ramping neurons and their average firing rate were significantly higher in context A compared with context B (Fig. 2e-f), suggesting that ramping amplitudes and rates preferentially encode the CS+ odor probability and not context or the odors sequence identity. This behaviorally relevant coding bias is consistent with the increase in the rate of responding neurons and their firing rates to the CS+ odor (Extended Data Fig. 5a-e), and with previous studies reporting enhanced representation of the behaviorally relevant stimuli ^11,32,33^. To further examine whether ramping amplitudes and rates encode the relative CS+ probabilities, we compared the ramping activity across the A, B, and M contexts. We found that the average amplitudes and rates of ramping neurons indeed scaled with the CS+ odor probability, that is, A > M > B (Extended Data Fig. 3d-f). This graded ramping activity emerged at the expert state (Fig. 2g) and suggests that ramping neurons preferentially encode the CS+ odor probability and not the context or the odor sequence identity. Ramping amplitude did not correlate with task reaction time across context (Extended Data Fig. 3h), suggesting that it does not reflect trial engagement. Furthermore, the differences in ramping amplitude in A and B contexts could not be attributed to differences in sniffing frequency (Extended Data Fig. 3i-j). Notably, ramping activity dropped sharply at the onset of the CS odors, regardless of the context (Fig. 2c and Extended Data Fig. 3g), indicating that ramping neurons unlikely reflect predictive sensory responses of the expected stimuli, as reported in other sensory cortices^9,10^, motor licking activity, or the stimulus-associated value. Similar, albeit weaker, learning-dependent enhancements in representation were also observed in neural responses to the context odors themselves (Extended Data Fig. 4), suggesting that probability-modulated changes in neural activity emerge as soon as mice identify the odor defining the context. Taken together, aPC ramping activity scales with the CS+ odor probability. This scaling disappeared as soon as the CS odors are perceived, because at that point, there is no ambiguity about odor identity.

**Fig. 2:**
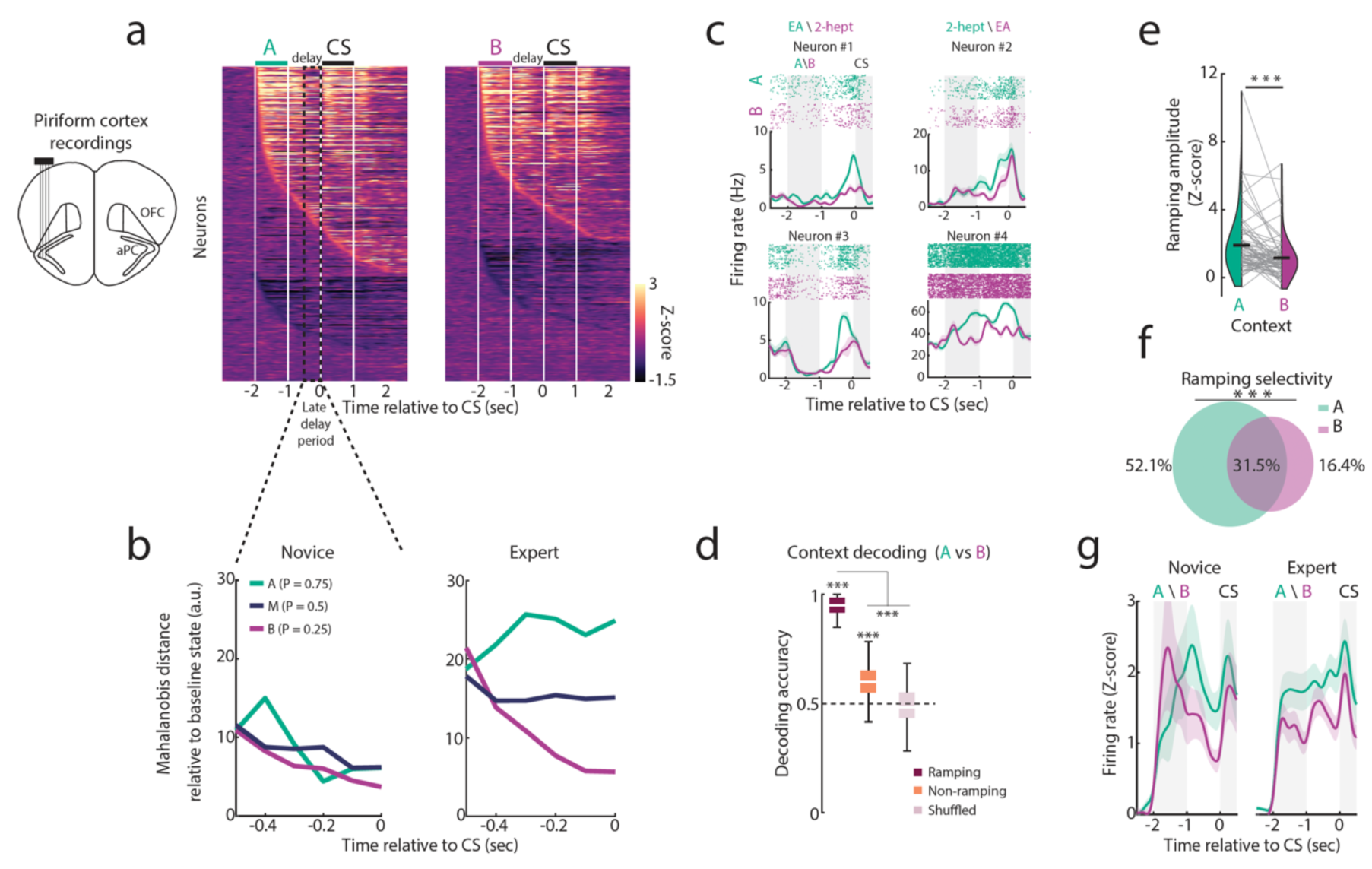
Ramping activity in aPC encodes the estimated probability of the behaviorally relevant odor. a. Two PSTH matrices showing all the recorded neurons during task performance in the A (left) and B (right) contexts (N = 401 neurons from four mice). PSTHs were Z-scored and sorted by peak latency of excitatory and inhibitory separately. For each neuron, all trials were pooled (both CS+ and CS-). White lines denote the duration of the context odors followed by the CS odor. Dashed black lines represent the late delay period used for subsequent analyses. b. Probability-modulated population trajectories in the neural state space following learning. The population activity of all recorded neurons was projected onto the first two principal components during the late delay period, separately for each context and learning state. In each time bin (100 ms), the distance of the trajectory was measured relative to the baseline state (time window -4 to -3, before trial onset). Values close to zero denote higher similarity to the baseline activity state. c. Four examples of ramping neurons, showing higher amplitudes in the A odor compared with the B odor. As the identity of context odor A and B were switched across mice, in neurons #1 and #3, odor A was ethyl acetate (EA) and odor B was 2-heptanone, while in neurons #2 and #4 the identities were reversed. d. Ramping neurons encode the trial context during the delay period. Context decoding analysis using the activity of ramping and non-ramping neurons (red and orange, respectively). A linear SVM decoder discriminated between the A and B contexts using the neural activity before the CS odor onset (-330 to 0 ms, Methods). The same analysis using shuffled neural activity is shown in mauve (P < 0.001, two-sample bootstrap relative to shuffled values and between ramping and non-ramping neurons). e. The mean population ramping firing rate in the A and B contexts, quantified in the time window before the CS odor onset (-330 to 0 ms; P < 0.001, two-tailed paired *t*-test). The black line marks the mean. f. A Venn diagram showing the relative percentage of neurons with significant ramping in A context, B context, and both (ramping selectivity, N = 73 ramping neurons). Significantly higher rate of ramping neurons was found in context A compared with context B (P < 0.001, χ^2^ goodness of fit to uniform distribution). g. The mean Z-scored PSTHs of ramping neurons recorded in novice and expert states (N = 12 and 61 ramping neurons, respectively).

### Odor-selective prediction-error signals in aPC emerge with learning

Our paradigm enables us to examine responses to the CS odors when they are predicted at different probabilities, and when these predictions are violated. According to the predictive-coding framework, the response elicited by an expected stimulus in the sensory cortex should be lower than that to a less expected stimulus ^22, 34^. Analyzing all CS odor responding neurons (N = 293/401 neurons; CS+, 256; CS-, 192; both, 155 neurons), we identified a subpopulation of neurons whose odor-evoked responses were modulated by the trial context (Fig. 3a-b; N = 48/293, 16% of all CS responding neurons). We termed this group ‘context-modulated’ neurons and the remaining neurons ‘sensory’ (i.e., context-independent response to the CS+, CS-, or both odors). Concordantly, a linear decoder successfully decoded the trial context identity from the neural activity of these context-modulated neurons, whereas it performed at chance level when applied to ‘sensory’ neurons (Fig. 3c). Context-modulated neurons are either sensitive to the context identity (e.g., the odor sequence in a specific context) or signal deviation from the prediction based on the learned contingencies. If context-modulated neurons encode context identity, one would expect to find similar average firing rates of context-preferring neurons for the A and B contexts (Fig. 3d, left). However, if these neuron responses are affected by the difference between the predicted and the actual odor, one expects an overall firing rates reduction when odors are highly predicted compared to when they are less predicted (Fig. 3d, right), a PE-like response. We found that the firing rates of most context-modulated neurons were significantly lower when the CS odors were highly predicted compared to when they were less predicted (Fig. 3e), demonstrating that a subpopulation of aPC neurons exhibits a PE-like odor response. This effect was observed for both CS+ and CS- odors, and primarily in the expert state, indicating that this modulated activity emerges with learning and is not restricted to the behaviorally relevant odor (Fig. 3e-f).

To examine how context-modulated neurons are affected by the magnitude of the error, we analyzed the context-modulated neurons when the error magnitude was 50% (i.e., context A versus B) or 25% (i.e., context M versus A or B). Interestingly, the change in firing rate did not scale with prediction level, as would be expected for canonical PE neurons. In contrast, the fraction of context-modulated neurons increased with the error magnitude (Extended Data Fig. 5f–g). These results suggest that PE-like signals in aPC might be represented at the population level, rather than through graded responses of individual neurons, as observed for dopaminergic PE signals. Interestingly, almost none of the neurons that responded to both CS odors were context-modulated for both odors (N = 2/155; Fig. 3b, g, and Extended Data Fig. 5h), indicating that these neurons are feature-selective and do not signal a general prediction violation.

In summary, we found that during learning of odor probabilities, two neural subpopulations emerged, which encode odor probabilities differentially: 1) ramping activity that carries information on the behaviorally relevant odor probability before its onset, and 2) context-modulated neurons that signal a deviation from prediction when the odor is detected, regardless of its behavioral relevance, and in an odor-selective manner. These two subpopulations have relatively little overlap, suggesting they subserved two distinct circuits (Fig. 3h).

### Prediction hampers odor representation

How different levels of prediction affect stimulus separability in the sensory cortex is debated ^5,8,10,15,20,35,36^. To examine this, we trained an SVM decoder using the activity of context-modulated neurons to discriminate between the CS odors (identity decoding) when they were both predicted with high or low probabilities. While the overall decoding rates were high for both prediction levels, the discrimination rate was significantly lower for the more predicted stimulus (Extended Data Fig. 5i, left bars), suggesting that reduced activity due to expectation hampers the stimuli representation. The decoding accuracy of the less predicted stimuli was similar to that achieved when using the ‘sensory’ neurons (Extended Data Fig. 5i, right bars), suggesting that, at least in our setup, prediction violation does not enhance odor representation.

**Fig. 3:**
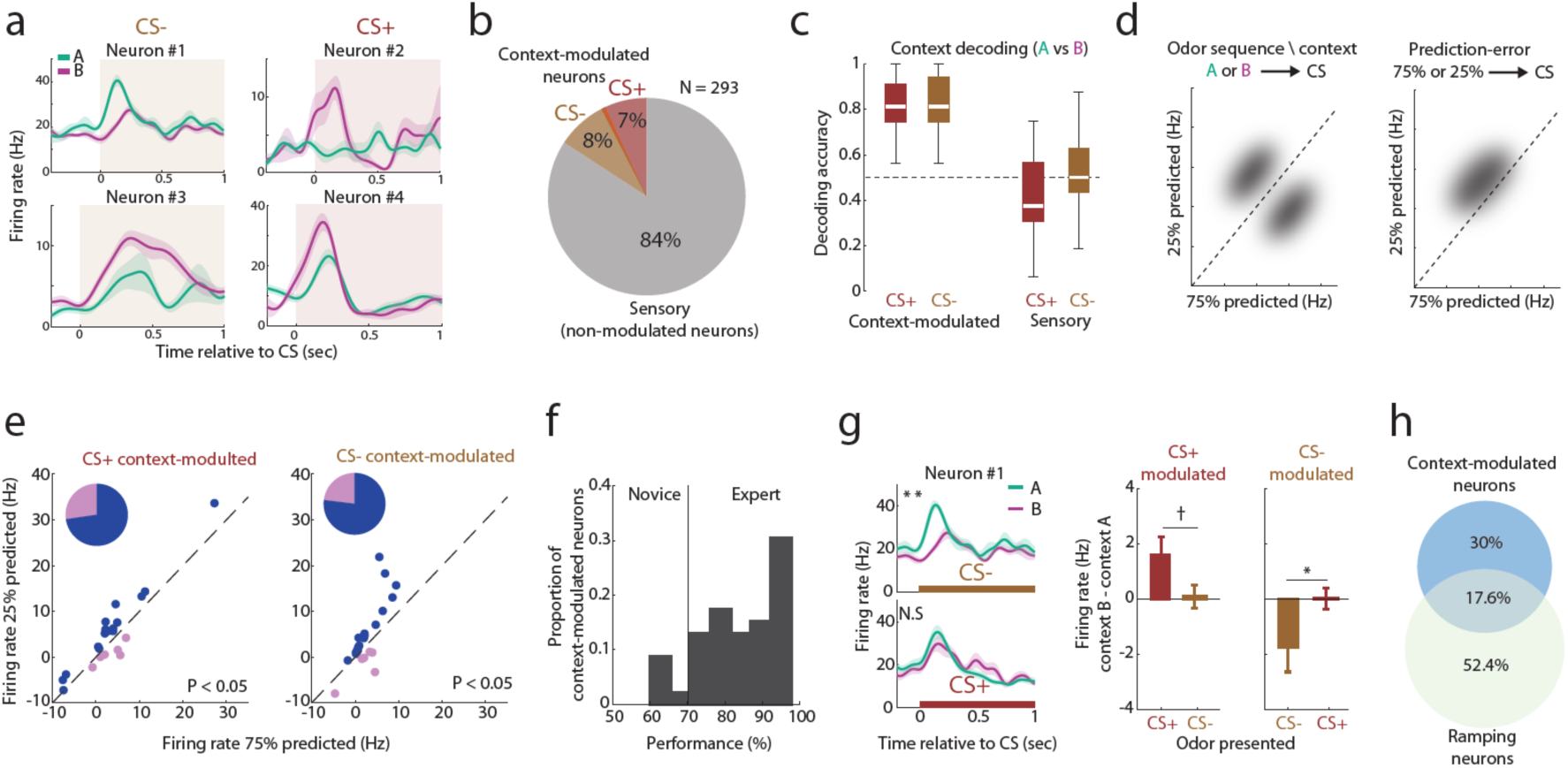
Odor-selective prediction-error signals in aPC emerge with learning. a. Four example neurons showing context-modulated responses to the CS- odor (neuron #1 and #3) or the CS+ odor (neuron #2 and #4). Neurons #1, #3, and #4 exhibit higher firing rates when the odor was unpredicted compared to the predicted response, while neuron #3 shows the opposite relation. b. A pie chart showing the percentage of context-modulated neurons for CS+ (red), CS- (brown), and both (overlap) out of all the CS odors responding neurons. Non-modulated neurons are considered ‘sensory’ (gray). c. Context-modulated neurons encode the trial context during CS odor responses. An SVM decoder was trained to discriminate between the odor-evoked activity in the A or B contexts (context decoding). Decoding accuracy is shown for CS+ and CS-odors in boxplots for context-modulated neurons (left) and for ‘sensory’ neurons (right). Decoder performance using the activity of ‘sensory’ neurons was at chance level. d. Two hypotheses about what information context-modulated neurons signal. Left: Context modulation is sensitive to context identity, resulting in similar firing rate among context-preferring neurons. Right: Context-modulation neurons show a PE-like response, with reduced responses when the odor is expected compared to when it is less expected. e. Firing rates of context-modulated neurons for CS+ (left, N = 22 neurons) and CS-(right, N = 26 neurons). Most neurons reduced their activity when the odor was highly predicted compared to when it was less expected (blue dots, P < 0.05, two-tailed paired *t*-test for both). Insets pie charts show the percentage of neurons that decreased (blue) or increased (pink) their activity when the odor was predicted at 75% for the CS+ (left) and CS- odors (right). f. A histogram of the proportion of context-modulated neurons as a function of task performance, showing the emergence of these neurons during the late stage of the novice state and peaking at the expert state. g. Context-modulation is odor-selective. Left: an example neuron showing odor response for both CS+ and CS- odors, but context-modulation only to the CS- odor. Right: Mean ± SEM difference in firing rate between contexts (B-A) for all context-modulated neurons is shown for the CS+ and CS- odors. (†P = 0.056 and *P < 0.05, two-tailed paired *t*-test). h. A Venn diagram showing the overlap between ramping and CS context-modulated neurons (N = 138 ramping and context-modulated neurons combined).

### Probabilistic information emerges in OFC before it appears in aPC

The representation of stimulus probabilities and other task variables in a primary sensory cortex is likely to be mediated through top-down projections from higher brain regions. The orbitofrontal cortex (OFC) forms projections to sensory cortices ^25,27,37,38^, is known to encode value ^39–41^, forms context-dependent reward associations and encodes reward confidence ^24,42–48^. The OFC projects to the aPC ^28,29^, however, it is currently unknown what information these projections carry. We hypothesized that OFC projections to the aPC convey context-dependent behaviorally relevant probabilistic predictions. To test this hypothesis, we first examined how task contingencies are represented in the OFC. Recording the activity in the OFC during task-learning (Fig. 4a; N = 3 mice, 157 well-isolated units, 78% task-modulated neurons), we found similar task-contingency representations to those found in aPC, in the form of ramping activity and context-modulated CS odor responses (Fig. 4b). As in aPC, ramping selectivity was biased towards the A context, and context-modulated neurons were highly odor-selective (Fig. 4b-c, e). Interestingly, we found that population activity in the OFC encode the CS+ odor probabilities during the delay already in the novice state, as evident from comparing OFC activity in the different contexts using population trajectories (Fig. 4d), the ramping neuron amplitudes and the rate of ramping neurons (Extended Data Fig. 6a). This could not be explained by different responses to the context odors as they did not differ in the naïve state (Extended Data Fig. 6e). These encodings emerged in aPC only in the expert state (Fig. 2b). By contrast, and similarly to the aPC, context-modulated neurons emerged in the OFC only after mice reached expert state (Fig. 4e), were odor-selective and exhibited a PE-like firing rate suppression (Fig. 4e and Extended Data Fig. 6b). The proportion of context-modulated neurons in the OFC was higher than in aPC, as might be expected from a frontal brain region (26% compared with 16% of all CS odor-responsive neurons, P = 0.015, propotion test). Moreover, as with the aPC, OFC neurons exhibited enhanced representation of behaviorally relevant odors during learning, as previously shown (Roesch et al., 2007; Schoenbaum and Eichenbaum, 1995). However, while in aPC, learning increased the CS+ odor-evoked firing rates and the percentage of CS+ odor-responsive neurons relative to the CS- odor (Extended Data Fig. 5a-e), in the OFC, learning suppressed both these measures for the CS- odor without affecting the CS+ odor (Extended Data Fig. 6c-d). Interestingly, these differences in odor responses between the two regions were also evident in the responses to the context odors (compare Extended Data Fig. 6e to Extended Data Fig. 4c-d).

To conclude, we found highly similar task-contingency representations in both brain regions. OFC population activity encoded odor probability already in the novice state, before mice showed pronounced differences in reaction times, preceding the changes in aPC neural activity. Moreover, the two brain regions exhibited distinct plasticity mechanisms in representing the behaviorally relevant odors throughout learning. We next asked how OFC activity affects task learning and performance.

### The OFC is required for learning and retrieving odor probabilities

Previous studies have shown that OFC silencing does not impair the learning of stimulus discrimination tasks, but hampers reward-related rule updating ^24,27,42^. To test how OFC activity affects the learning of probabilistic contingencies, we expressed the inhibitory DREADD receptors (AAV8-hSyn-hM4Di) or a control virus (AAV8-hSyn-mCherry) bilaterally in the OFC (Extended Data Fig. 6f). To silence OFC activity during task performance, we injected the JHU37160 dihydrochloride (JH60), a high-potency DREADD agonist (Bonaventura et al., 2019), before each session. Verification of JH60 effect showed robust suppression of OFC activity ∼20-30 minutes post-injection, as reported previously ^51,52^(Extended Data Fig. 6g-h). All JH60-injected mice learned to discriminate between CS+ and CS- at a similar rate to mice with intact OFC (Fig. 4f and Extended Data Fig. 6i, N = 5 mice, 43 sessions). This confirms that learning odor-reward associations is not conspicuously affected by OFC suppression.

Strikingly, lick reaction times, a proxy for learning the odor probability contingencies (Fig. 1), did not differ between contexts (Fig. 4g and Extended Data Fig. 6k), indicating that OFC suppression impaired learning of odor probability contingencies. Mice eventually learned the probabilities after reaching very high task performance (Fig. 4g). The lick reaction times in the B condition were not different between mice with and without OFC suppression, suggesting that OFC silencing did not affect lick motivation (Extended Data Fig. 6l). We next examined whether the OFC is needed after mice have learned the probability contingencies. To test this, we repeated the above experiment in a new cohort of mice, where we suppressed OFC activity after mice had learned the probability contingencies (i.e., reached the expert state; Fig. 4h; N = 4 mice, 30 sessions). We found that expert mice’s ability to utilize probability information was hampered, as indicated by similar lick reaction times in the A and B contexts following OFC suppression (Fig. 4i-k and Extended Data Fig. 6m). Overall, these findings dissociate the role of the OFC in learning stimulus probabilities and odor-reward associations, revealing a previously unknown key role of the OFC in learning and retrieving stimulus probabilities.

**Fig. 4:**
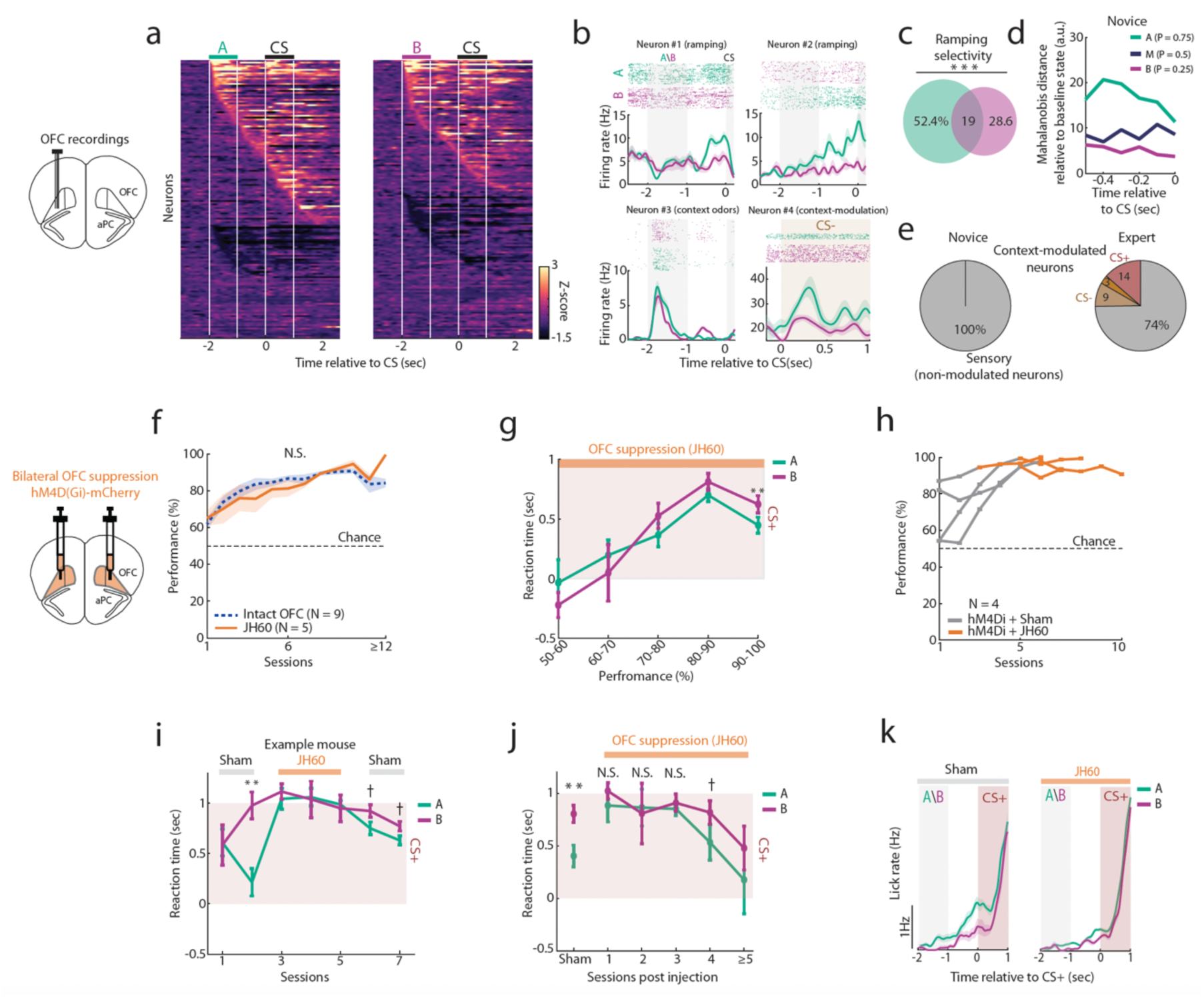
The orbitofrontal cortex is required for learning and retrieving odor probabilities. a. Two PSTH matrices displaying all the recorded OFC neurons in the A and B contexts (N = 157 neurons from three mice). b. Similar OFC neural dynamics to those found in aPC. Four example neurons showing ramping activity (neurons #1-2), a response to both A and B context odors (neuron #3), and a CS- odor context-modulation (neuron #4). c. A Venn diagram showing the ramping neuron selectivity (P < 0.001, χ^2^ goodness of fit to uniform distribution). d. Probability-modulated OFC population activity during the delay period emerges in the novice state. The distance in the state space between the population trajectories relative to the baseline during the late delay period (compare with Fig. 2b). e. Distribution of context-modulated and ‘sensory’ neurons in novice and expert states. Context-modulated neurons emerged only in the expert state and are odor-selective. f. Left panel: Schematic illustration depicting the bilateral injection of DREADDs (AAV8-hSyn-hM4Di-mCherry) to the OFC. Right panel: OFC suppression did not significantly affect odor discrimination learning. Shown are the mean ± SEM learning curves of all mice with OFC suppression (N = 5). For comparison, the average learning curve of mice with intact OFC is shown (dashed blue curve, N = 9 mice). No significant difference was found between mice’s task performance with and without OFC suppression in any session number. g. OFC suppression impairs the learning of task probabilities. An analysis of the lick reaction times in contexts A and B as a function of task performance (compare to Fig. 1g). Significantly shorter reaction times in context A emerged only when mice performed at 90-100% in the odor-discrimination task (**P < 0.01, two-tailed paired *t*-test). h. OFC suppression at the expert state did not affect task performance. All learning curves of mice that learn the probability contingencies (reached expert state) with sham injection (grey) and with OFC suppression (orange, JH60; N = 4 mice, 30 sessions). In sham sessions, mice either received saline (N = 2) or no injection (N = 2). i. Lick reaction times before, during, and post JH60 injection in an example mouse. Reaction times were not different between contexts A and B during sessions with JH60 injections, and they were shorter in context A compared to context B in sham sessions with saline (**P < 0.01 and †P < 0.1, two-tailed paired *t*-test, see Extended Data Fig. 6k for full lick PSTHs). j. Reaction times in contexts A and B before and after OFC suppression. The mean ± SEM reaction times are shown as a function of the session number post JH60 injection, and before JH60 injection (left, sham, P < 0.01 two-tailed paired *t*-test). Reaction times were similar between the two contexts in the first three sessions post-injection (N.S., P > 0.3, two-tailed paired *t*-test), and became different in the following sessions (†P = 0.055, two-tailed paired t-test). k. Average lick PSTH in contexts A and B in sham (left, N = 13 sessions) and OFC suppression sessions (JH60, right, N = 17 sessions), showing reduced anticipatory lick rate in context A following OFC suppression.

### Probability, but not context information in aPC requires the OFC

The failure to exploit the probability information when the OFC is suppressed could result from impaired encoding of the context identities (Fig. 5a, hypothesis #1). Alternatively, context information could still be available (i.e., neurons that discriminate between A and B contexts), but the probability values were either not correctly assigned to the contexts or lost (Fig. 5a, hypothesis #2). To differentiate between these two hypotheses, we recorded aPC activity during task performance while suppressing OFC activity throughout the learning process (Fig. 5b and Extended Data Fig. 7a-b; N = 3 mice, out of the five mice reported in Fig. 4, N = 425 well-isolated units; 70% task-modulated neurons). We found that the average ramping activity no longer scaled with the CS+ probability (Fig. 2b-e), resulting in similar average ramping amplitudes in contexts A and B, which resembled the ramping amplitudes of the B context recorded in mice with intact OFC (Fig. 5c-d). Such degraded probability representation was also observed in the population activity in the neural state space (Fig. 5e). Interestingly, the rate of ramping neurons was no longer biased towards the A context, indicating that the information about the estimated CS+ probability is impaired (Fig. 5f and Extended Data Fig. 7d; compare to Fig. 2f). Notably, most ramping neurons remained selective to one of the contexts (Fig. 5f), suggesting that context information could still be extracted, as verified by successful decoding of the trial context (Fig. 5g; context decoding). The rate of context-modulated neurons was similar to the rate found in mice with intact OFC (N = 43/236, 17% of all CS responding neurons, N = 28 and N = 15 for CS+ and CS-, respectively, Fig. 5h), further indicating that context information is still available in aPC even when the OFC is silenced. Importantly, whereas in mice with intact OFC, context-modulated neurons reduced their firing rate when the odors were highly predicted compared to when they were less predicted, when OFC is silenced this bias disapeared (Fig. 5i), such that there was an equal rate of neurons that increased or decreased their firing rate (54%-46%, see the left hypothesis in Fig. 3d). Interestingly, this effect was observed only in the CS+ odor responses (Extended Data Fig. 7e), which is consistent with the selective reduction in CS+, but not CS-, odor-evoked activity following OFC suppression (Extended Data Fig. 8a) and the reported role of the OFC in value coding. This equality in the rate was also observed when we applied this analysis to the ‘sensory’ neurons or to neural data with no contextual prior (passive odor stimulation, Extended Data Fig. 7f-g, Methods). As in the case of ramping neurons, an SVM decoder could still reliably decode the context identity from the activity of the context-modulated neurons, indicating that context information is still available (Extended Data Fig. 7h-i). These findings suggest that the information about the probability of the behaviorally relevant odor, but not the context identity, is impaired in aPC following OFC silencing. Finally, the enhanced representation of the behaviorally relevant odors (i.e., CS+ and A odors) that emerged in aPC with learning was abolished following OFC suppression (Extended Data Fig. 8, compare with Extended Data Fig. 4 and 5).

To summarize, we revealed an OFC-sensory cortex circuit that is required to encode probability information in aPC for behaviorally relevant odors and is necessary for guiding appropriate probability-driven behavior.

**Fig. 5:**
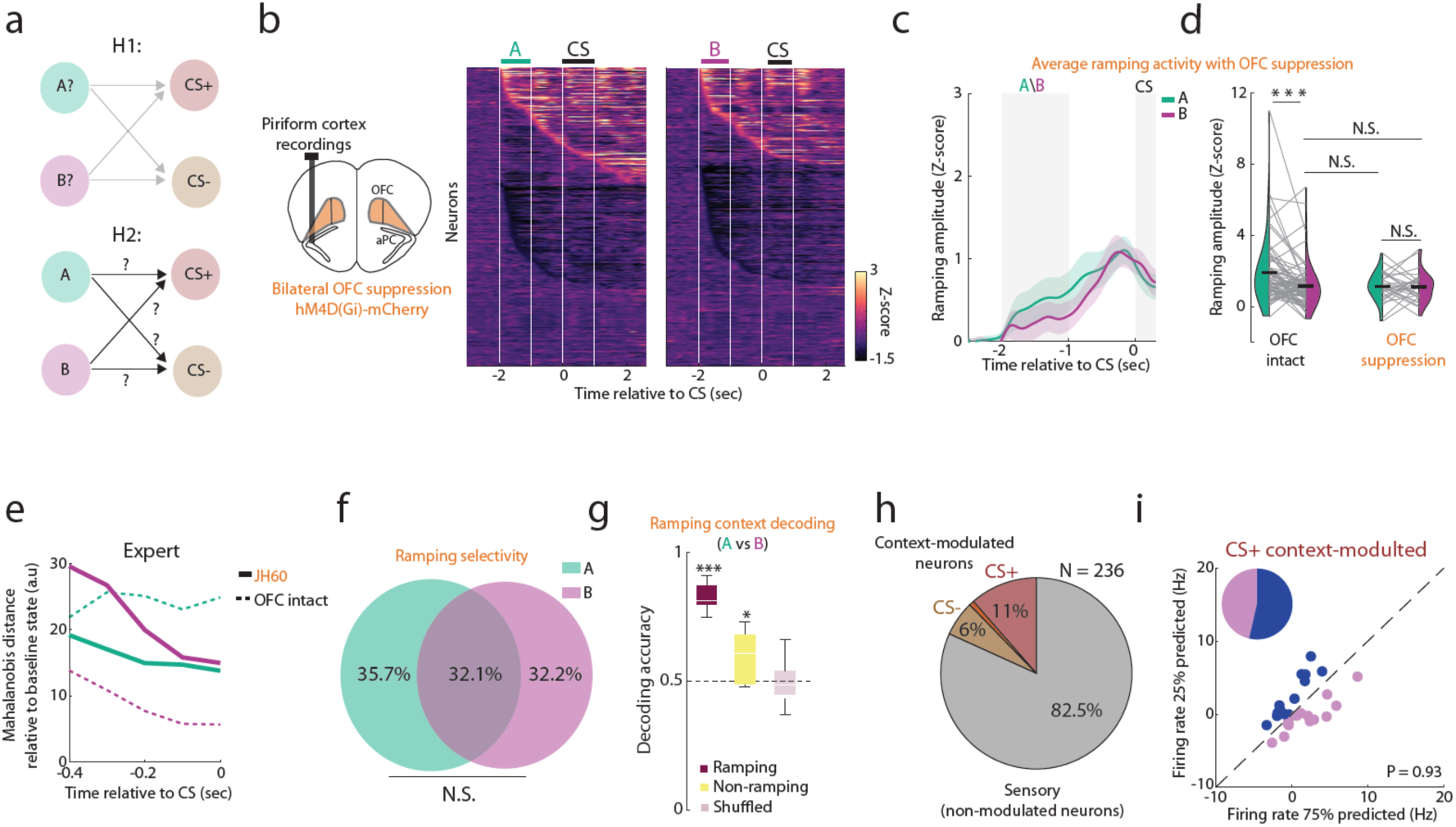
Probability, but not context information in aPC requires the OFC. a. Two hypotheses on the effect of OFC activity on aPC probability coding. In the first hypothesis (H1), silencing OFC impairs the encoding of contexts in aPC, whereas in the second hypothesis (H2), silencing OFC does not impair context encoding, but rather the probabilities are not correctly assigned to the contexts. The data support the second hypothesis. b. Two PSTH matrices showing all aPC neurons recorded during OFC silencing (N = 425 neurons from three mice). c. The average (Z-scored) activity from all ramping neurons. No significant difference in ramping amplitude was found on average between A and B contexts (P = 0.94, two-tailed paired *t*-test). d. Z-scored ramping amplitudes recorded in mice with intact (left, P < 0.001, two-tailed paired *t*-test) and suppressed (right, P = 0.94, two-tailed paired *t*-test) OFC. Ramping amplitude in the A context is similar to the level of the B context that was recorded in mice with intact OFC. e. OFC silencing impairs probability-modulated population trajectory in the neural state space. Similar to Fig. 2b, population activity in expert sessions was projected onto the first two principal components during the late delay period, and the distance of the trajectory was measured relative to the baseline state (time window -4 to -3, before trial onset). For reference, the distances obtained in mice with intact OFC (Fig. 2b) are shown in dashed lines. f. Nearly uniform ramping selectivity distribution following OFC suppression. A Venn diagram showing the overall selectivity of ramping neurons. Ramping selectivity distribution was not significantly different from a uniform distribution (P = 0.88, χ^2^ goodness of fit to uniform distribution). g. Context decoding analysis using the activity of ramping and non-ramping neurons during OFC suppression (red and yellow, respectively). An SVM decoder discriminated between the A and B contexts using neural activity during the late delay period (shown in the inset, -330 to 0 ms; Methods). The same analysis using shuffled neural activity is shown in a mauve box plot (*P < 0.05, ***P < 0.001, two-sample bootstrap relative to shuffled values). h. A pie chart showing the percentage of context-modulated neurons in CS+ (red) and CS- (brown) responsive aPC neurons during OFC suppression. Non-modulated ‘sensory’ neurons are colored in gray. i. Firing rates of CS+ odor context-modulated neurons (N = 28) were predicted in high or low probability. No difference was found between the two conditions (P = 0.93, two-tailed paired *t*-test). The number of neurons that reduced (blue) or increased (pink) their response when the CS+ odor is similar. See Fig. 3e for comparison with intact OFC recordings, where aPC neurons reduced their firing rates following high-probability prediction.

## Discussion

In this study, we revealed an OFC-aPC neural circuit involved in learning and retrieving odor statistics in rapidly changing context-dependent behavioral settings. Mice were engaged in a task in which odors had different probabilities of occurring depending on the context (Fig. 1). To evaluate micès internal probability model as learning progressed, we tracked lick reaction times and false alarm rates and found that mice inferred and utilized the latent context-dependent odor statistics to enhance their behavior. At the expert level, mice still exhibited faster lick reaction times in the context of the higher CS+ probability but delayed their reaction time until after the presentation of the CS odors, suggesting that mice also learned the state transition model. This task design enabled us to investigate how single neurons and population activity in the aPC and OFC encode rapidly changing context-dependent odor probabilities across different learning stages. We found two distinct neuronal subpopulations that emerged with learning and encoded odor probabilities differentially before and during the predicted odor event. We further identified a previously unknown role of the OFC in learning and retrieving stimulus probabilities, and in propagating this probability information to the olfactory cortex.

A subset of aPC and OFC neurons ramped up their activity during the delay period up to the CS odor onset. Ramping amplitudes scaled with the CS+ probability (i.e., A > M > B contexts), suggesting ramping neurons take part in a circuit that preferentially encodes the estimated probability of the behaviorally relevant odor. This scaling emerged with the learning of task contingencies, and was observed regardless of the odor identity that served as the A context odor (Fig. 2d). These findings indicate that ramping neurons are unlikely to encode the identity of the odor, context or the odor sequence, but rather the learned contingencies. Ramping activity has been previously observed during decision-making and goal-directed behavior in higher brain regions ^4,41,53,54^ as well as in primary sensory cortices ^9–11^, where it was suggested to anticipate the timing of stimulus occurrence or its identity. Here, we show that ramping activity also encodes the estimated probability of upcoming stimulus occurrence. These findings are in contrast to the ramping activity of midbrain dopaminergic neurons, which is positively correlated with the reward uncertainty ^1,2,55^. At the CS odor onset, ramping neurons typically reduced their activity to baseline, regardless of the actual odor identity (Extended Data Fig. 3g), indicating that ramping activity is neither an early predictive response to the upcoming odor nor a reflection of motor licking activity. This reduction in activity further suggests that ramping neurons do not encode the expected value, since it differs between CS+ and CS- odors. Once the CS odor is detected, the probability information becomes irrelevant, as reflected in a reduction in neuronal activity.

While ramping amplitudes were modulated by the probability values before CS odor onset, a distinct subpopulation exhibited probability-modulated responses to the CS odors, with neurons reducing their firing rates when the odor was highly predicted compared to when it was less predicted suggesting that these signals odor PE. Several recent studies reported stimulus PE-like signals in the neocortex, ^5,9,12,15–17,19,35,36^, however, there is no evidence for PE signals in the olfactory cortex, a three-layer paleocortex. Furthermore, some of the above studies manipulated the stimulus prediction level by increasing the stimulus’s presentation rate, making it difficult to dissociate expectation from habituation and/or familiarity. To account for these possible confounds, all stimuli in our design were presented for a similar number of trials in a pseudo-random order. Thus, our results reveal the existence of PE-like signals even when all stimuli are equally familiar and temporally predictable, indicating that they signal violations of probabilistic estimation.

In classical frameworks, PE corresponds to a signed mismatch between actual and predicted inputs and is typically expressed as a graded, bidirectional signal at the level of individual neurons. However, studies examining mismatch signals in the sensory cortex have typically failed to identify clear bidirectional responses in individual neurons. This has led to the proposal that PE in sensory areas may be implemented through distinct neuronal populations, with separate neurons encoding positive (more-than-expected) and negative (less-than-expected) errors ^22^. Consistent with this view, a small population of neurons responding to omitted stimuli has been reported in a sensory cortex ^9, 15^. In contrast, the extent to which sensory neurons exhibit graded scaling of responses with prediction level has been less systematically explored, although a recent work suggests such scaling may occur in the auditory systems of songbirds ^8^.

In our study, neurons did not exhibit graded scaling with prediction level at the individual-neuron level. Instead, the degree of deviation from expectation was reflected as a graded change in the fraction of modulated neurons (Extended Data Fig. 5f-g), suggesting a population-level representation of this key feature in PE frameworks. Future studies are required to test the hypothesis that PE in cortical circuits may be conveyed through population recruitment rather than through graded mismatch computations within individual neurons.

In the visual cortex, PE signals were evident in neurons highly selective for the stimulus^15^. While pyramidal neurons in the visual cortex are typically narrowly tuned, aPC neurons can have relatively wide tuning curves ^56–59^, making it an attractive brain region to test stimulus-selective PE in neurons that are less selective. Interestingly, we found that aPC neurons that reliably responded to both CS odors were modulated by the prediction level of only one odor. These finding points suggest these signals are more aligned with a PE-like computation rather than a general expectation-suppression, and importantly suggest to a flexible mechanism by which neurons encode predictions in a circuit-selective manner. This implies that signaling a prediction violation is not a feature that defines a neuron type, but rather depends on the odor identity. Future research is required to study these circuit-dependent signals, their extension to multiple odor identities or reward associations. That said, we found similar rates of neurons modulated by predictions for rewarded and unrewarded odors, indicating that these signals do not depend on the expected value. This may suggest that these neurons signal a ‘state prediction error’ of the learned statistical model, that is, reporting the discrepancy between the predicted and observed stimuli, regardless of their reward associations ^60^.

How different levels of prediction affect stimulus separability in the sensory cortex is debated. Several recent studies showed improved representation of unexpected stimuli ^15,20,36^, while others suggested that expected stimuli are better represented ^5,8,10,35^. We found that a high prediction level of both odors degrades their separability in the neural space compared to when they were less predicted. However, the overall representation of these odors is unlikely to be strongly affected, as the rate of neurons signaling prediction error is relatively low (16% of all CS odor-responding neurons, similar to a recent report ^19^) and the majority of the neurons robustly represent the odor identity regardless of its prediction level (i.e., “sensory” neurons). This low rate of neurons signaling prediction error is consistent with the idea that their role is to signal deviations from the current estimate to downstream neurons and therefore do not contribute to stimulus representation ^22^.

A recent study demonstrated that, in the visual cortex, stimulus prediction errors signal saliency in the face of a novel stimulus rather then the difference between actual and expected stimulus responses ^15^. We recorded PE-like signals when both CS odors are equally familiar and temporally predicted, suggesting that PE may serve additional roles. aPC PE-like signals emerged at the expert level, scaled with the PE magnitude (Extended Data Fig. 5), and importantly were attenuated upon OFC silencing, impairing the mouse’s ability to utilize probability information. This suggests that PE signals play an additional key role in the formation of internal statistical models that represent the estimated context-dependent stimulus probabilities.

Overall, our data suggest that PE signals, sensory neurons, and ramping neurons in aPC are components of distinct neural circuits serving distinct roles: reporting deviations from a learned model, representing stimulus identity, and encoding stimulus occurrence probability, respectively.

While earlier studies showed that the OFC is required for reward-related behavior ^25,26^, more recent studies have shown that the OFC is not essential for learning basic stimulus-reward associations ^24,27^. Consistent with this, we found that mice could learn and retrieve the CS-reward associations when the OFC was inactive. Notably, mice failed to learn the odor probabilities or utilize previously learned probability information when the OFC was silenced. These findings point to a key, previously unknown role of the OFC in representing probabilistic state transitions, consistent with the growing evidence that the OFC is involved in constructing a cognitive map of the task ^47,49,52,61–63^. The findings that probability and context information emerge in the aPC and depend on the OFC, may suggest that features of this cognitive map propagate to sensory regions.

The OFC forms reciprocal projections with the aPC ^28,29^, however, the functional role of these projections is largely unknown. Our data revealed similar task-related activity signatures in the aPC and OFC, including ramping activity and odor-selective PE neurons. We found that the OFC encoded task probabilities during the delay period already at the novice state, whereas this representation emerged in aPC at the expert state, and that this representation in aPC was abolished following OFC silencing. These findings suggest that the OFC is the source of probability information in aPC. Notably, OFC silencing hampered CS+ but not CS- odor PE signals. Taken together, this suggests that during task learning, the OFC modulates aPC activity (directly or indirectly), to represent the relative probabilities of the behaviorally relevant odors. Consistent with ^50^, we found that as task learning progressed, both aPC and OFC exhibited enhanced representation of the behaviorally relevant odors. However, we further demonstrated here that these two regions achieved this via distinct plasticity mechanisms: aPC neurons increased their responses to reward-associated odors, while the OFC neurons reduced their responses to non-rewarded odors (Extended Data Fig. 4–6). Interestingly, silencing the OFC did not affect context representation in aPC, indicating that context information in aPC is maintained independently of OFC input. We hypothesize that context information could arise through recurrent activity within the aPC ^57,64^ or through context representation in the hippocampal formation^24,65^ In Summary, we demonstrate that context-dependent mapping of event probabilities in the sensory cortex arises from interactions with the frontal cortex.

## Acknowledgments

This study was supported by a grant from the Israel Science Foundation program [1211/25].

## Author Contributions

T.D. performed the experiments and analyzed the data, T.D. and R.H. conceptualized the experiments and wrote the paper.

## Methods

All surgical and experimental procedures were conducted in accordance with the National Institutes of Health Guide for the Care and Use of Laboratory Animals and Bar Ilan University guidelines for the use and care of laboratory animals in research, and were approved and supervised by the Institutional Animal Care and Use Committee (IACUC). Mice were housed in a room on a reverse light/dark cycle. Before surgery, mice were housed in a group cage of up to five mice and received no experimental treatment, except genotyping. After surgery, mice were housed individually. All experiments were performed during the dark period. Twenty C57BL male mice (Envigo) aged 3-12 months were used in this study.

### Surgical Procedures

Mice were briefly anesthetized with isoflurane before receiving an intraperitoneal injection of ketamine/medetomidine (60/0.5 mg/kg i.p.), and then fixed in a stereotaxic frame. Body temperature was maintained at 36-37°C using a homoeothermic blanket system (Harvard Apparatus). The skin above the dorsal skull was removed, and a headplate was implemented using dental cement (Parkell). For tetrodes implementation, a small craniotomy was made for the implementation of the ground screw. Another craniotomy of ∼0.4mm in diameter was made for the insertion of the tetrodes (aPC: A-P, 1.8-2.2mm, M-L, 2.4-2.6mm; OFC: A-P, 2.2-2.4mm, M-L, 1.5-1.8mm). Tetrodes were lowered slowly to the region of interest (D-V: aPC, -3.5mm; OFC, -3mm), and were fixed to the skull using dental cement, and a cape was built around for protection. The overall structure weighted ∼4 grams. An analgesic injection (Carprofen) was administered at the end of the surgery, and for three days every 24 hours. Mice were allowed to recover for at least one week before behavioral training started.

### Viral Injection

Mice were briefly anesthetized with isoflurane before an i.p. injection of ketamine/ medetomidine (60/0.5 mg/kg i.p.), and then fixed in a stereotaxic frame. AAV8-hSyn-hM4Di-mCherry (titer: 1.43x10^13^ particles/ml) or AAV1-hSyn-mCherry (titer: 7×10¹²; University of North Carolina Gene Therapy Center) were injected bilaterally into the orbitofrontal cortex. The coordinates were ±1.25 mm (M-L) from the midline rhinal fissure, +2.4-2.8 mm (A-P), and -1.8-2.5mm (D-V), 200 nL in each hemisphere. Viruses were injected using a syringe (Hamilton Company) attached to a micro-injector (IMS-10, Narishige, Japan) at a rate of 70 nl/ min, which was left in place for 5 minutes to allow viral particle diffusion before needle removal. Incisions were closed with tissue glue (Vetbond), and an analgesic injection (Carprofen) was administered at the end of the surgery, and for three days every 24 hours. Mice were allowed to recover for at least 4 weeks before behavioral training and electrophysiology were carried out.

### Behavioral setup

The behavioral setup was built inside a noise-proof Faraday cage (med associations). Head-fixed mice were placed on a custom-built round treadmill. Task control and data acquisition were implemented using custom LabView (NI) scripts. Lick detection was achieved by placing an IR sensor between the lick port and the mouse, such that each beam crossing (lick) triggered a voltage change. Water solenoids (The Lee Company) triggered 4-8uL of sucrose (5%) in each rewarded trial. Respiration signal was measured using a custom pressure sensor that was placed nearby the nostril to detect pressure changes (Honeywell). Coned-shaped odor port was placed ∼3 cm away from the animal, and all odor pipes converged into the tip of the port. Photoionization detector (PID, Aurora Scientific) measurements were carried out before the training of each animal started.

### Odors Application

Odors were applied using a custom-built olfactometer. Odors were diluted in mineral oil (1:100) and stored in sealed plastic tubes. This concentration was chosen to elicit a detectable response. The odors were refreshed before each session (5-10ml). Airflow was controlled with a mass flow controller (Agilent, Alimc-2LSPM) and set to 0.3 slpm. Air circulated freely between stimulations to reduce odor remnants. A vent placed behind the mouse removed residual odors. All odors used were from Sigma-Aldrich at their highest purity. The odors were ethyl butyrate (CAS: 105-54-4), ethyl acetate (CAS: 141-78-6), 2-heptanone (CAS: 110-43-0), and pentyl acetate (CAS 628-63-7). Odors A and B were 2-heptanone or ethyl acetate (90 and 48 sessions with 2-heptanone or ethyl acetate as A, respectively), and the CS+ and CS- odors remained fixed across all mice (ethyl butyrate and pentyl acetate, respectively). Odor stimulation times and sequences were controlled by custom LabView scripts.

### Behavioral training

#### Water restriction

Mice were water restricted for two days before training began. And received 1ml of water per day. Behavioral sessions occurred once per day during the dark phase and lasted for approximately 50 minutes or until the mouse stopped performing, whichever came earlier. Mice would receive all their water from these sessions, unless it was necessary to supply additional water to maintain a stable body weight (>85% of initial body weight). Water intake was monitored daily, and on days without training, mice received 1 ml of water.

#### Pretraining

Before training started, mice were handled, head-fixed in the setup, and were given drops of sucrose water freely from the lick port. This procedure habituated them to head fixation and allowed adjustment of the IR sensor placement so that licks are clearly detected. After habituation to the lick port, mice were presented with the CS+\-odors only, and reward was available in CS+ trials. This session was essentially an odor discrimination task without the predicting odors, in order to familiarize the mice with the CS+\ odors, and typically lasted 50-100 trials. In these sessions, success rates were typically at 50-60%, as mice still acclimatized to the lick port and to pair the CS+ odor with reward availability. Training with the experimental paradigm started after one to two pretraining sessions.

#### Probability training

Mice were presented in each trial with two temporally separated odors. The first, the context odor, could be either odor A or B, was applied for one second. 75% of the trials after odor A ended with CS+, and 75% of the trials following odor B ended with CS-. We verified that changes in neuronal activity could not be attributed to such bias in the number of trial in each condition. Following the context odor there was a delay period of one second without odor stimulation, and after this period either the CS+ or CS- odors were presented for one second. In most sessions sucrose reward was given after a decision window of one second after the CS+ odor offset, but in some sessions the window length was 500ms or 0ms. Inter-trial-interval ranged between 8-15 seconds across sessions. The number of trials starting with A or B odors and ending with CS+ or CS- odors was similar. In two mice we introduced another context odor, M, which predicted the CS+\- odors at equal probabilities (50%). In these sessions the total number of trials was divided by three. To control for effects of odor preference, the identity of context odors A and B was switched across mice. No punishment was given for false alarms. A Hit trial is defined as a trial with at least three licks in the time duration between the CS+ onset and the reward onset. To accelerate learning, reward was delivered at the end of the decision window even if the mouse did not perform the sufficient number of licks. The number of trials per session was 144 ± 35, mean ± standard deviation; N = 138 sessions across all mice. Discrimination performance dropped to chance level when expert mice performed a session with no odor application, indicating task performance did not depend on other sensory cues (N = 2 mice).

#### Probability reversal

in seven mice, after mice reached expert level (>70% performance) and had significant difference in lick reaction times between A and B contexts, we introduced probability reversal (data not shown). Following reversal, the identity of the A and B odors was switched while the identity of the CS+\- odors remained fixed. Probability reversal was not cued or signaled to the animal, started only from the beginning of a session, and were no longer switched back in the following sessions. Since the CS+\- identity was not switched, there was no drop in the discrimination task accuracy, however, reactions times were affected. Mice were typically accustomed to the new probabilities after one session, and some mice showed rapid switch by exhibiting shorter reaction times in the new A context after ∼50-100 trials.

### Electrophysiology

The extracellular spiking activity and the local-field potential (LFP) were recorded using custom-built microdrives with eight tetrodes. Neural signals were first filtered at 0.1–9,000 Hz and then at 600–6000 Hz to obtain spiking activity. Signals were sampled and recorded at 32 kHz (Cheetah data acquisition software, Neuralynx). Tetrodes were chronically implemented in the left anterior piriform cortex, and were lowered ∼80-100 um at the end of each recording day to sample an independent neurons population across recordings sessions. Tetrodes placement at each session was estimated by depth and confirmed by histology. Overall we recorded 401 neurons in the aPC with OFC intact, 157 neurons in the OFC and 425 neurons in the aPC with OFC silencing, yielding a total of 983 recorded neurons.

### Chemogenetic manipulation

Before each session, mice (N = 5) were briefly anesthetized in isoflurane (4% in air) and received an i.p. injection of JHU37160 dihydrochloride (JH60; 0.1 mg/kg, dissolved in PBS). Mice were left in their home cage for 20 min, to allow the DREADD agonist to effectively inhibit hM4di transfected OFC neurons. In another cohort of mice (N = 4), OFC suppression was introduced after mice reach expert level. In the sessions prior to the expert level mice were briefly anesthetized 20 minutes before training and were given either an i.p. injection of PBS or no injection (sham). Electrophysiology recording was carried out to verify the effect of JH60 injection on hM4Di infected OFC neurons. Recording was done using 32-sites silicone probe (Neuronexus) in an anesthetized mouse four weeks post viral injection. Recoding started at the time of JH60 injection, and odors were applied throughout the recording to verify neuronal suppression at the present of sensory stimuli (Extended Data Fig. 6).

### Histology

To assess tetrodes trajectories and viral expression, after experiment completion the brain was removed and was fixed in 4% PFA for at least 24 hours. The tissue was then suspended in 3% agarose in PBS. A vibratome (Leica, VT1200) cut coronal sections of 60-80 µm that were mounted and subsequently imaged with a fluorescence microscope.

### General statistical analysis

All data analyses were performed in MATLAB. The number of data points used for all statistical tests and graphs are detailed in the Fig. or Fig. legends. Significance was defined as a p-value of 0.05 or 0.01 for all tests. Unless stated otherwise, we report the mean ± SEM or %95 confidence interval. Mean ± standard deviation is reported when estimated from a bootstrap process. We use statistical tests as required by the test’s null hypothesis and population assumptions, and all tests were two-tailed. We did not use statistical methods to predetermine sample sizes.

### Data analysis

#### Spike sorting

Spike signals were sorted and clustered offline using Spike3D (Neuralynx). Only visually well-isolated clusters were used, with less than 5% of spikes violating an inter-spike interval of 3ms (median violation of refractory period across all neurons is 0.75%).

#### Behavioral analysis

Respiration signals were band-passed filter at 0.1-20Hz. The onsets of inhalation and exhalation were defined as the zero-crossings of the signal that came before and after the peak, respectively. Lick signals were converted to binary vectors using threshold crossing. A low-pass filter of 12.5Hz was applied to allow maximal inter-lick-interval of 80ms. Odor events were registered using a TTL channel. Hit or false alarms were assigned if there were at least three licks between the CS odor onset and the end of the decision window (typically one second), which was followed by reward delivery in CS+ trials. Reactions times were registered as the timestamp of the first lick in the lick bout, after the trial onset (time point -2 second relative to CS odor onset). If lick bout started before trial onset and continued into the start of the trial, reaction time was corrected to the first lick following the onset of the CS odor. In the rare cases of lick threshold crossing following paws activity or non-discriminative constant licking, a trial was discarded if the IR signal was above threshold in more than 45% of the trial duration. Sniff rasterplots were constructed using the inhalations onsets, marking the start of a respiration cycle. Both lick and sniff rasterplots were aligned to the CS odor onset. Lick and sniff PSTHS were constructed by convolving the spike rates using a Gaussian window with 70ms standard deviation. Success rates were computed for each session as Hit-(1-FA)/2, where Hit and FA are the fraction of trials with licking activity out of all CS+ and CS- trials, respectively.

#### Odor-evoked responses

An odor-evoked firing rate was computed over the odor duration, subtracted by the activity during a two-second baseline window before trial initiation. A response was defined as significant if the mean spike rate during the odor duration was significantly different from the activity during baseline (P < 0.01, two-tailed paired *t*-test). Responses were sorted to excitatory and inhibitory if the mean spike rate during odor presentation increased or decreased relative to baseline, respectively. PSTHs were smoothed using a Gaussian window with 70 or 100ms standard deviation, and were z-scored using the activity during a baseline period before trial onset. For CS+\- odors, we defined a significant response by pooling all trials from A and B contexts. Neurons were sorted into novice (<70%) and expert (>70%) groups based on the odor discrimination success rate obtained in the sessions they were recorded at. Quantification of task modulated neurons was done for A and B contexts separately during the context odors duration (-2 to -1 sec), during the late delay period (-.33 to 0 sec) and during the CS+ and CS- presentation. A task modulated neuron had significant activity change during one of the above time windows or contexts relative to baseline.

#### Population PCA trajectories

PCA analysis was performed over non-overlapping time bins of 100ms to generate population PCA trajectories during the delay period. For each context and learning state, the Mahalanobis distance was computed using the first two principal components between the trajectory in the i^th^ time bin and the mean trajectory values during the baseline activity (time bins -4 to -3).

#### Hierarchical clustering

Z-scored PSTHs of all neurons in the A context were used for the clustering analysis, which used a Pearson correlation as the distance function and was restricted by a maximal number of 14 clusters. The clusters indices were then applied to the B context z-score PSTHs, and the mean cluster activity was plotted for both contexts.

#### Analysis of neuronal subpopulations activity

*Ramping neurons* the mean firing rate during a time period of 330ms to 0 was compared to the mean firing rate during baseline, two seconds before trial initiation. Neurons with mean firing rate higher than two standard errors of baseline activity and evoke firing rate above 2 Hz were classified as ramping neurons. This time window was based on previous studies that analyzed neural activity during a late delay period ^9,10,31^.

*Sniff to ramping correlation* for each ramping neuron from a session with valid recording of respiratory activity, we computed the correlation between the activity during the 330ms time window prior CS onset across trials and the sniff rate during this time window.

*Context modulated neurons:* for each neuron, the mean activity during the CS+\-odors (1 seconds) was compared between the A and B conditions. Neurons with significantly different activity for at least one of the CS odors (P < 0.05 two-tailed paired *t*-test) were classified as context-modulated neurons. We quantified the percentage of context-modulated neurons relative to the total number of neurons that responded to each CS odor (P < 0.05, one-sample proportion test against ɑ = 0.05).

#### Response modulation to CS+\- in passive odor stimulations

For each neuron that responded to CS+, CS- or both in passive stimulation (Extended Data Fig. 2), we randomly sampled 75% and 25% of the trials and compared the means of the two samples. We repeated this process for each neuron 100 times, such that for each neuron we got ∼5% (false-discovery rate) of the repetitions that had significantly different activity between the 75% and the 25% samples of the trials (P < 0.05, two-tailed paired *t*-test). Within the ∼5% in each neuron we counted the number of cases where R(75%) > R(25%) and vice versa, where R denoted the mean firing rate across trials. We then computed the distribution of neurons that increased or decreased their activity.

#### Classification analysis

*Passive odor stimulation:* a support-vector machine (SVM) classifier with 5-fold cross-validation was used to classify the odor identity from the neural activity recorded during passive odor stimulation. A sliding window of 200ms was used for the moving classification analysis.

*Context decoding:* the activity of context-modulated neurons during a one-second period of odor presentation was used to predict the prior odor context (A or B) using a binary SVM classifier with 5-fold cross-validation. The same analysis was performed over non-context-modulated (‘sensory’) neurons. The same analysis was done over a time window of -330ms to 0, to decode the context from the activity of ramping neurons.

*CS+\- identity decoding in high or low prediction* CS+ to CS- firing rates during a one second period of odor duration were used. Trials were sorted for 75% prediction, that is R(CS+|A) and R(CS-|B), and 25 prediction R(CS+|B) and R(CS-|A). The number of trials was balanced. Classification using SVM 5-fold cross-validation classifier was performed using trials.

## Extended Data Figures

**Extended Data Fig. 1:**
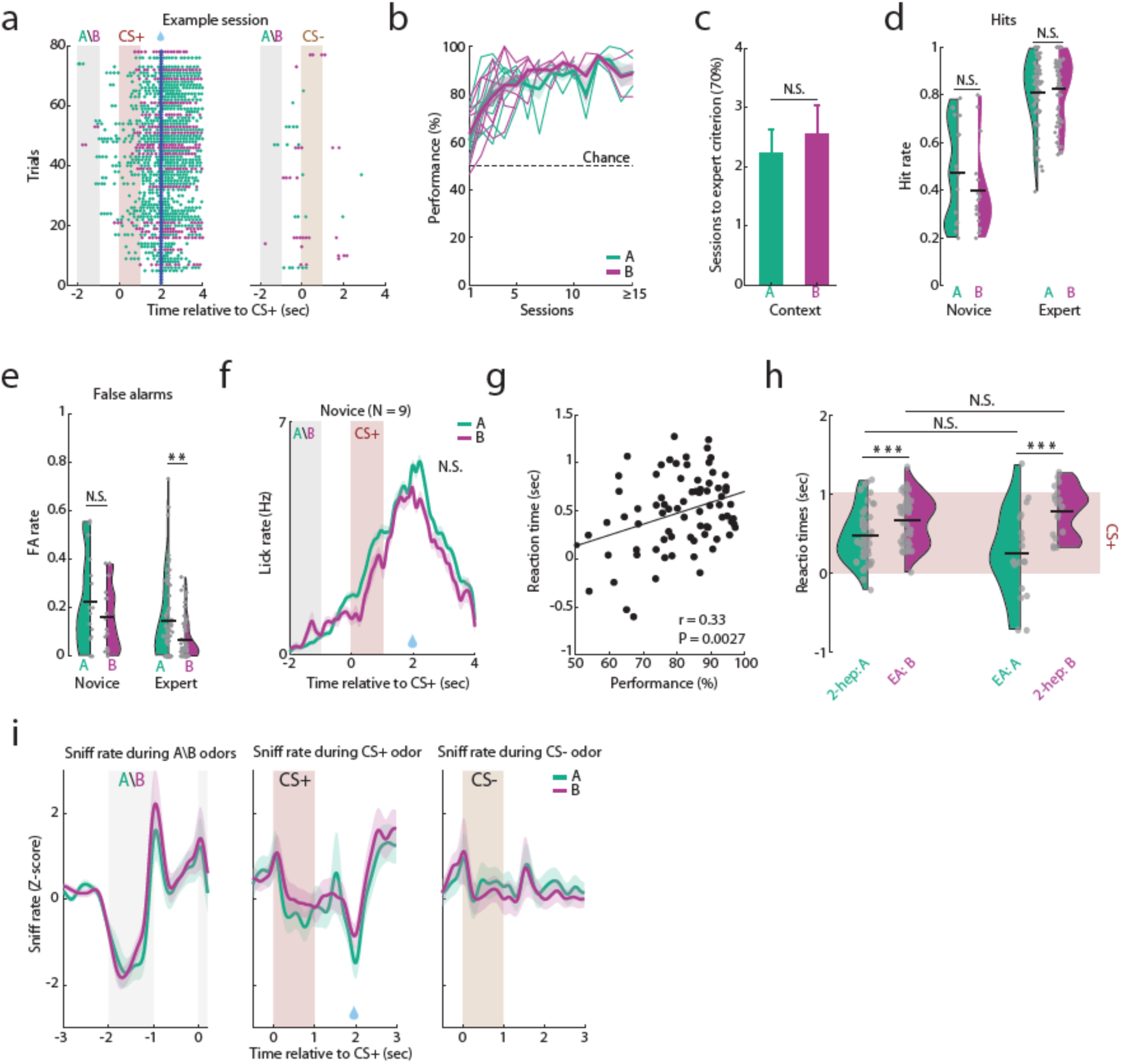
Additional analyses of the behavioral task. a. Two lick raster plots of an example session recorded in an expert mouse split into the CS+ (left) and CS- (right) odors. Green and purple represent trials of A or B contexts, respectively. Blue dot marks reward delivery. b. Learning curves of the nine mice, divided into context A and B trials. Mean ± SEM is shown in a bold line. Thin lines denote individual mice. c. No difference in learning rate between contexts A and B. Mean ± SEM of the number of sessions required to reach the 70% criterion (P = 0.47, N = 9 mice, two-tailed paired *t*-test; range 1-5 sessions). d. Hit rates in novice and expert mice across contexts A and B (N.S., P > 0.05, two-tailed paired *t*-test). e. Same as d, for false alarm rates (N.S., P = 0.27; **P < 0.01, two-tailed paired *t*-test) f. Licks PSTH in novice sessions, showing no significant difference in anticipatory licking rate between the two contexts (N.S., P = 0.32, two-tailed paired *t*-test). g. The relation between the mean reaction time and the mice’s performance in each session. Reaction times are delayed as learning progresses (Pearson correlation, r = 0.33, P = 0.0027, N = 78 sessions). h. Reaction times in the expert state are independent of the identities of the A and B odors. Reaction times are shown when odors A and B were 2-heptanone (2-hep) and ethyl acetate (EA) and vice versa (***P < 0.001 for both, two-tailed paired *t*-test). i. The average sniff rate (Z-scored) during the context odors (left), during the CS+ and CS- odors for contexts A and B (middle and right). (N = 9 mice across 70 sessions with valid respiration signal).

**Extended Data Fig. 2:**
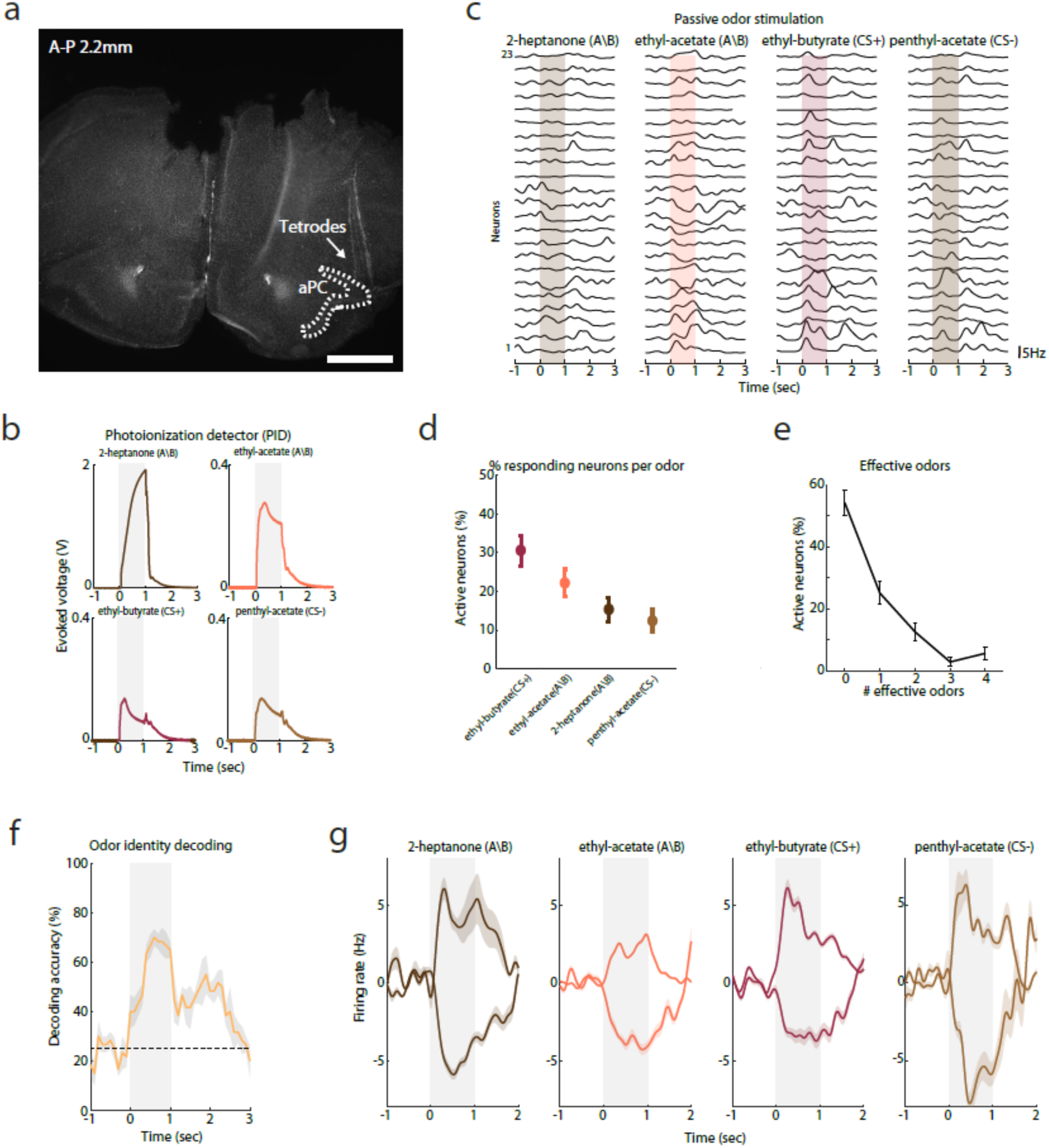
Passive odor stimulation before task learning. a. An example coronal section showing the trajectory of the tetrodes towards the aPC (marked with white dashed lines). A-P is 2.2 mm from Bregma, scale bar, 1 mm. b. Photoionization detector (PID) measurements of the odor output for the four odors used in the experiment. c. An example of 23 simultaneously recorded neurons from a session of passive odor stimulation, showing the average PSTH across trials for each neuron. The two left odors served as the context odors, and the two right odors were the CS+ and CS-, respectively. d. The percentage of neurons that responded to each odor (N = 6 mice, 72 neurons). e. Percentage of neurons activated by different numbers of odors. f. SVM classification of odor identities plotted against the odor duration. Chance level (25%) is shown with a dashed line. g. Average PSTHs of all the neurons that responded in excitation or inhibition to each odor.

**Extended Data Fig. 3:**
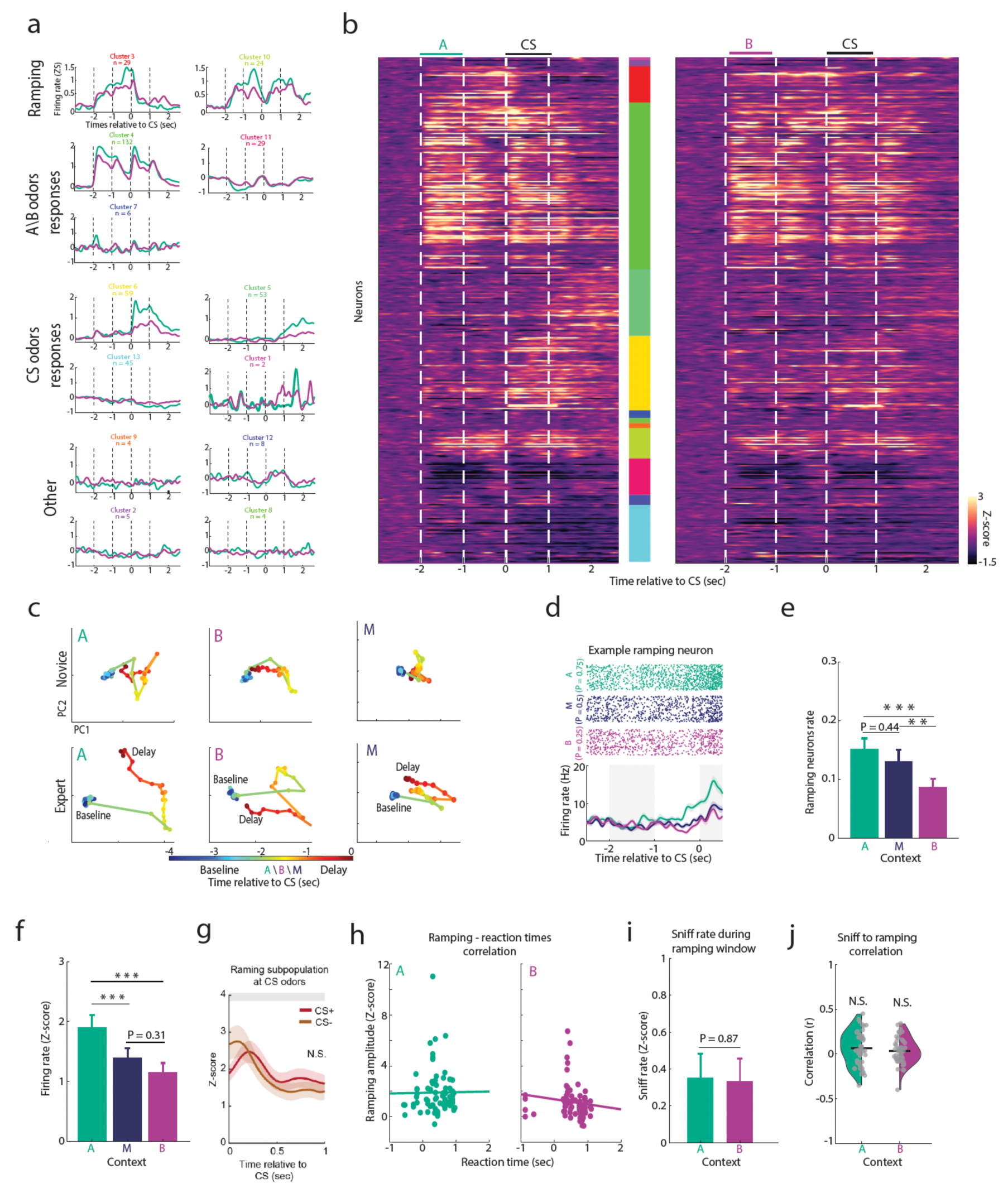
Additional analyses of ramping neurons’ activity. a. Hierarchical clustering of all recorded neurons. Clustering was performed on the PSTHs in the A context, and the cluster indices were then applied to the B context. Different clusters describe context-dependent activity during the context odor duration, the late delay period (ramping), and during the CS odor responses. b. Two PSTH matrices are sorted by the cluster identities. Color-coding on the right corresponds to the cluster numbers shown in **a**. c. Population trajectories of aPC activity as a function of the trial time. Population activity was projected onto the first two principal components separately for A, B, and M contexts, as well as novice and expert states (upper and lower rows, respectively). The time is color-coded, with each point reflecting 100 ms. The delay period was used for the distance analysis shown in **2b**. d. An example of a ramping neuron recorded in a mouse trained with three contexts, A (green), M (blue), and B (purple). e. The percentage of ramping neurons for the three contexts out of the total number of task-modulated neurons (***P < 0.001, **P < 0.01, two-tailed paired *t*-test). The percentages scale with the relative probability of CS+ odor occurrence. f. Ramping amplitudes scale with the CS+ probability. The mean ± SEM Z-scored ramping amplitude for A, M, and B (N = 73 neurons in A and B contexts and 61 neurons in M condition; ***P < 0.001, two-tailed paired *t*-test). g. Ramping activity collapses similarly at the CS+ and CS- onset. The mean activity of all ramping neurons during CS+ and CS- presentation. h. Ramping activity does not correlate with reaction times. For each context, the ramping amplitude of each neuron was plotted against the average reaction time obtained in that session (A context, r = 0.01, P = 0.92; B context, r = -0.13, P = 0.25, Pearson correlation). i. Ramping amplitudes cannot be explained by changes in sniffing activity. The mean ± SEM Z-scored sniff rate during the ramping analysis window (-330ms to 0) is shown for A and B contexts (P = 0.87, two-tailed paired *t*-test). j. Correlation between ramping and sniff activity. For each neuron, we computed the correlation between the activity during the 330 ms before the CS odor onset and the sniff rate in A and B contexts (P > 0.05, two-tailed one-sample *t*-test, see Methods).

**Extended Data Fig. 4:**
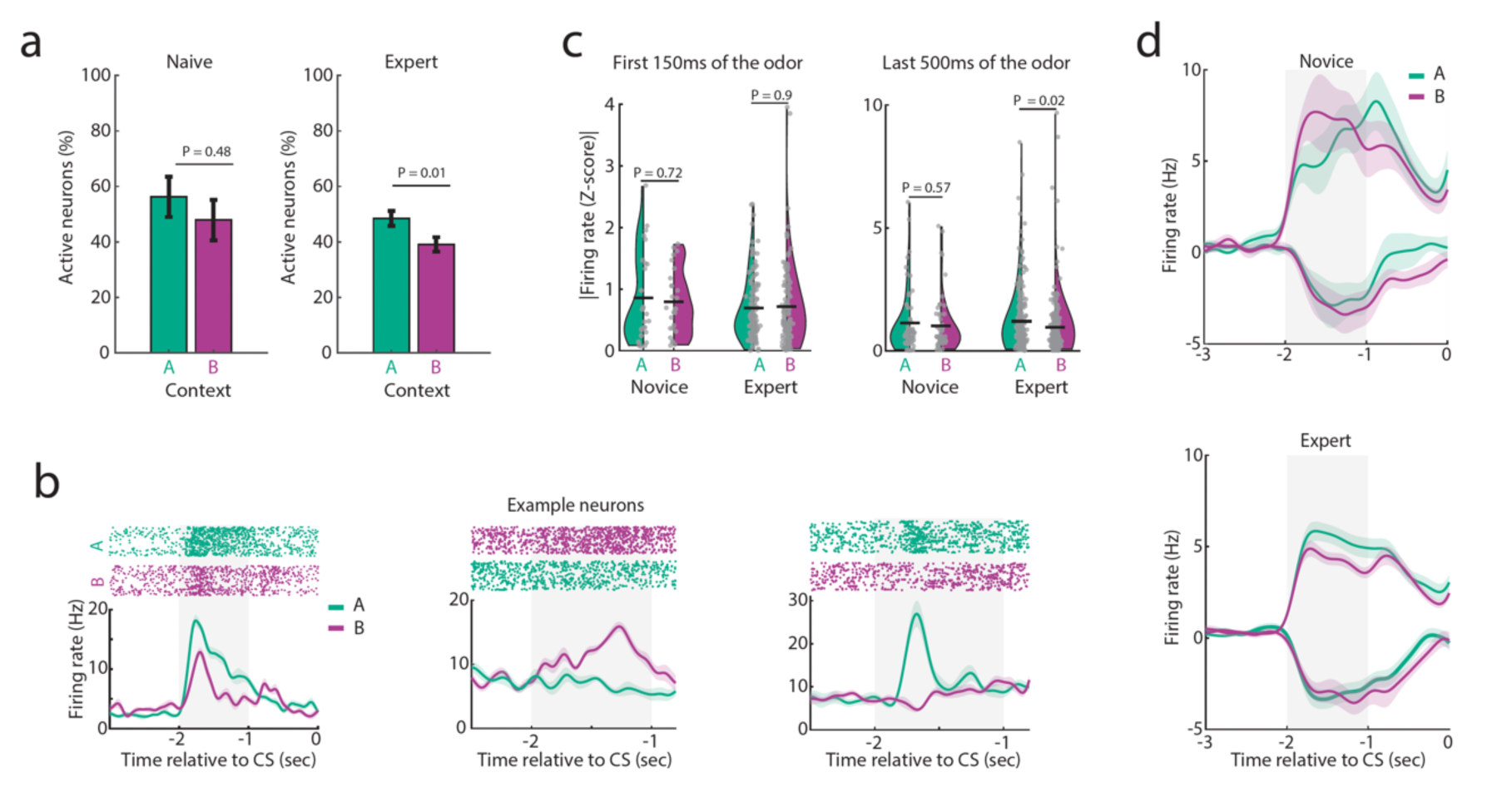
Analyses of aPC responses to the context odors. a. Quantification of context-odor responsive neurons in novice and expert states. Significantly more neurons were active during the A odor compared with the B odor in the expert state, but not in the novice state. See Extended Data Fig. 2 for quantification of the rate of neurons responding to these odors during passive stimulation. b. Three example neurons showing modulated response amplitude to the A and B odors in the expert state. c. Summary and statistics of the evoked activity for the A and B odors in novice and expert states. Data is shown for the first 150 ms of the odors (left) and for the last 500 ms of the odors (right). In the first 150 ms, there was no significant difference in firing rate between the two odors, as expected from a pure sensory response. Over the last 500 ms, odor A exhibited a significantly higher firing rate than odor B (P = 0.02, two-tailed rank sum test), reflecting the change in information content provided by the different contexts. d. The average PSTH of all context-odor responding neurons for A and B contexts, in novice (upper) and expert (lower) learning states. Higher response amplitude to the A context odor emerged with learning.

**Extended Data Fig. 5:**
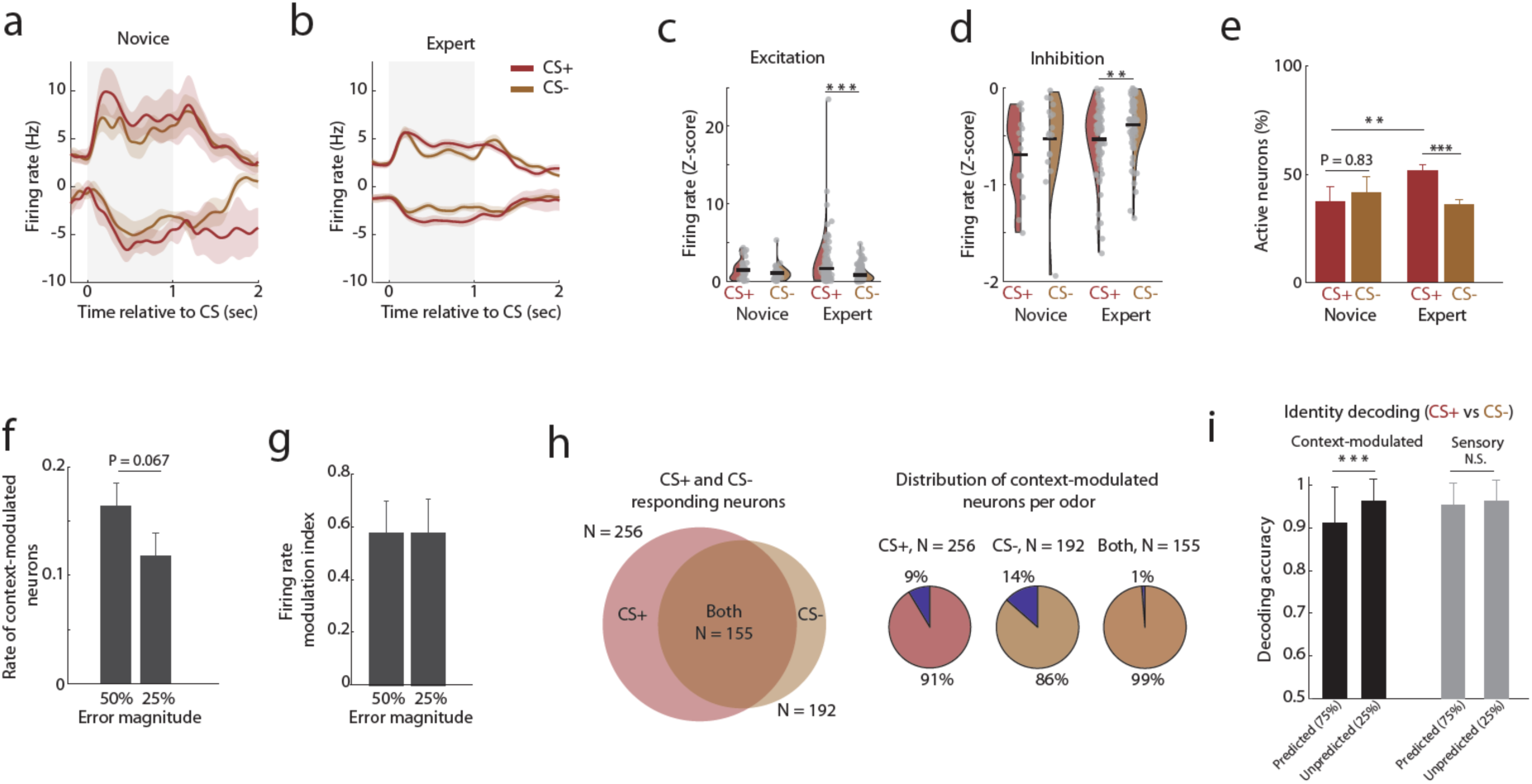
Learning-dependent changes of CS+ odor representation and additional analyses of context-modulated neurons. a. The average PSTH of all neurons that responded to CS+, CS-, or both, and were recorded in a novice (left). b. The same applies to neurons recorded in an expert state. c. Learning-dependent increase of CS+ activity. Statistical quantification of the data shown in **a-b**, for excitatory (left) odor responses. d. Same as **c**, for inhibitory odor responses. e. Percentage of active neurons in response to CS+ and CS-, recorded in novice and expert states. Significantly more neurons responded to the CS+ odor in the expert but not novice state (P = 0.83 and ***P = 3.0^-5^ for novice and expert, respectively; two-sample proportion test). The percentage of CS+ responding neurons increased with learning (novice versus expert, **P < 0.01, two-sample proportion test). f. The rate of PE signals scales with the magnitude of the error. The rate of context-modulated neurons is shown when the prediction difference was 50% (context A vs. B) and 25% (context M vs. A or B; P = 0.067, two-sample proportion test). Data is pooled for CS+ and CS- odors. g. Similar firing rate change in PE signals for different error magnitudes. Shown is the absolute firing rate modulation index pooled for CS+ and CS- odors, when the prediction difference is 50% and 25%. h. Left: a Venn diagram showing the number of CS+, CS-, and both odors responding neurons. Right: pie-charts showing the percentage of context-modulated neurons found for CS+, CS-, and both, relative to the number of neurons that responded to each odor or both (shown above each pie chart). i. Prediction degrades odor representation. An SVM decoder trained to discriminate between the CS+ and CS- odor-evoked responses (odor identity decoding), when they were predicted at high (75%) or low (25%) probabilities. Accuracy rates degrade when the odors are highly predicted (P < 0.001, two-sample bootstrap). No difference in classification rate was found when using the activity of ‘sensory’ neurons.

**Extended Data Fig. 6:**
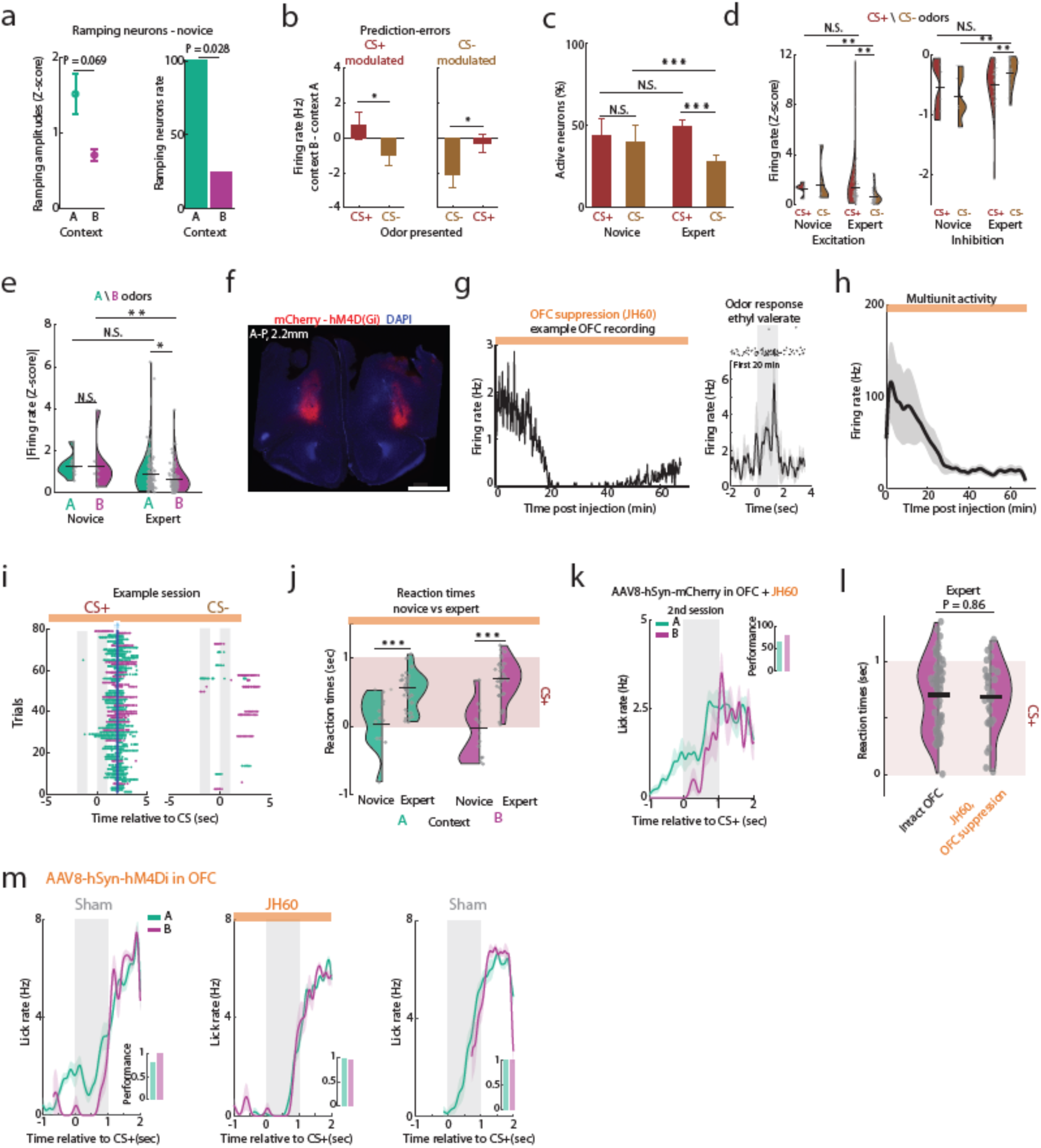
Additional analyses of OFC recordings and mice task performance during OFC suppression. a. Left: OFC ramping amplitudes in novice state (P = 0.069, two-tailed paired *t*-test). Right: Ramping neurons rate in the novice state (P = 0.028, two-sample proportion test). b. Context-modulated neurons in OFC are odor-selective and signal the odor prediction error. Mean ± SEM difference in firing rate between contexts (B-A) for all context-modulated neurons is shown for the CS+ and CS- odors (CS+: P = 0.012, CS-: P = 0.028, two-tailed paired *t*-test). c. Percentage of OFC responding neurons to the CS+ and CS- odors in novice and expert states. The percentage of CS- responding neurons was significantly reduced relative to the CS+ responding neurons in the expert state and the CS- responding neurons in the novice state (***P < 0.001, two-sample proportions test). d. OFC odor-evoked firing rates (Z-scored) of CS+ and CS- odors in novice and expert states, divided for excitatory and inhibitory odor responses. CS- odor responses were reduced compared to CS+ in the expert state (**P < 0.01, two-tailed paired *t*-test). e. Firing rate analysis (Z-scored) of all context odor-responsive neurons recorded in the OFC. Excitatory and inhibitory responses were pooled for statistical power. In the expert state, odor-evoked activity to the B context odor was reduced relative to the A odor and relative to B odor in the novice state (*P < 0.05 and **P < 0.01, two-tailed paired *t*-test). f. A coronal section from an example mouse showing bilateral expression of hM4Di in the OFC overlaid over DAPI (red and blue, respectively). Scale bar, 1mm. A-P distance is shown relative to Bregma. g. OFC suppression validation. Left: The firing rate of an example OFC neuron is plotted as a function of the JH60 injection time. Twenty minutes after the injection, the firing rate of the neurons was significantly suppressed. A typical session duration was ∼50 minutes (pink). Odors were applied throughout the recording to verify that suppression not only occurred in baseline conditions. Right: raster plot and a PSTH of the same neuron, showing a response to ethyl valerate during the first 20 minutes. The neuron was suppressed even during odor simulation in later trials. h. The average effect of JH60 injection on OFC activity. The mean ± SEM multiunit activity from all the electrode contacts located within the OFC. i. An example session from a trained mouse showing lick raster plots for the CS+ and CS- odors. Green and purple denote A and B contexts, respectively. Blue dots mark the reward. j. Reaction times were delayed in both contexts as learning progressed (P < 0.001, two-tailed paired *t*-test). The orange bar represents OFC suppression, and the red patch indicates the duration of the CS+ odor. k. An example session from a mouse injected with a control virus (AAV8-hSyn-mCherry), where JH60 was applied before the session. A significant difference in reaction time was observed between the two contexts, showing intact learning of odor probabilities (P < 0.01, two-tailed paired *t*-test). l. OFC suppression does not affect lick motivation. Reaction times in the B context with and without OFC suppression were not significantly different (P = 0.86, two-tailed paired *t*-test), showing no delay in lick activity due to OFC suppression. m. Lick PSTHs from the example mouse shown in Fig. 4i. Lick PSTHs from sham sessions (left and right, grey) before and after a session with JH60 (middle, orange), showing difference in anticipatory lick rates during sham but not JH60 session. Insets show the task performance in each session for A and B contexts (green and purple bars, respectively).

**Extended Data Fig. 7:**
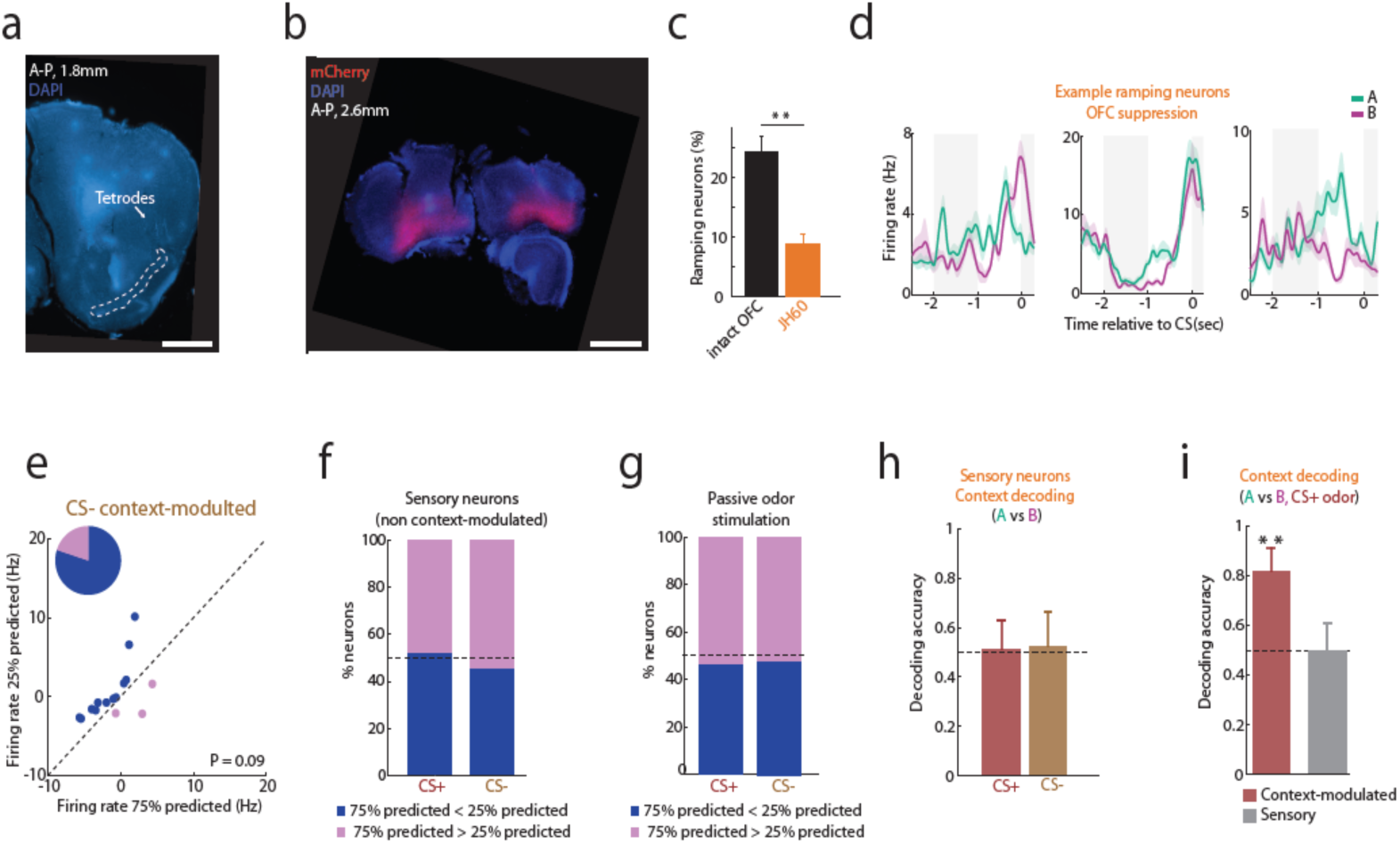
Additional analyses of ramping and context-modulated neurons in aPC during OFC suppression. a. A coronal section showing the tetrode’s trajectory (marked by white arrow) toward the aPC (marked by dashed white line) in a mouse with hM4Di-mCherry in the OFC. Scale bar, 1mm. b. Another more anterior coronal section of the same mouse, showing hM4Di-mCherry (red) in the OFC, overlaid over DAPI staining (blue). Scale bar, 1mm. c. Percentage of ramping neurons out of all task-modulated neurons, in mice with and without OFC suppression. d. Three examples of ramping neurons during OFC suppression. Examples show neurons that ramp for the B context (left), both contexts (middle), and the A context (right). e. Firing rates of CS- odor context-modulated neurons during OFC suppression (N = 15). Most neurons reduced their firing rates when the odor was highly predicted (blue, CS-|B). f. Percent of ‘sensory’ neurons that reduced (blue) or increased (pink) their firing rates when the odor (CS+ or CS-) was highly predicted. Analysis of all sensory neurons (N = 245 neurons) that did not exhibit significant context-modulated odor-evoked activity to at least one of the CS odors. The distribution is almost 50%-50% as expected by chance. g. Distribution of CS odor firing change without contextual prior, during passive stimulation. For each neuron that responded to CS+, CS-, or both, we randomly sampled 75% and 25% of the trials and compared the means of the two samples. We repeated this process for each neuron 100 times, such that for each neuron, we obtained ∼5% (false-discovery rate) of the repetitions that showed significantly different activity between the 75% and 25% samples of the trials (P < 0.05, two-tailed paired *t*-test). We then counted the number of significant repetitions in which the response of the 75% trials was higher than that of the 25% trials, and vice versa, to obtain an estimate of the type of error signal. h. Context decoding during CS+ presentation using the activity of ‘sensory’ neurons. Same analysis as in **Fig. 3c**. i. Probability decoding using the activity of CS+ context-modulated neurons. Same analysis as in **f**.

**Extended Data Fig. 8:**
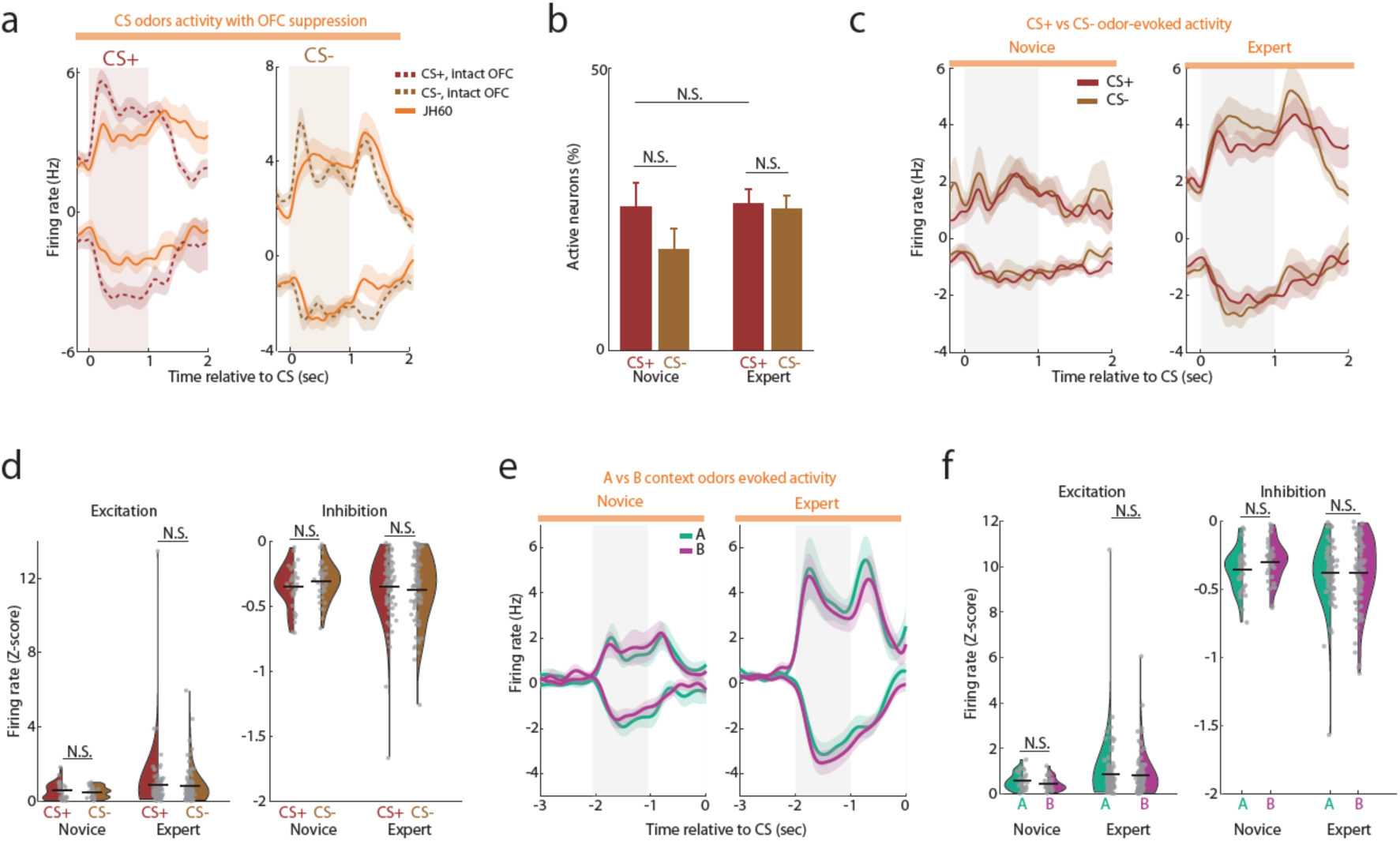
Additional analyses of aPC activity during OFC suppression. a. The average PSTH of all CS+ (left) and CS- (right) odor responding neurons, with and without OFC suppression (orange and dashed red or brown, respectively). OFC suppression reduced CS+ excitatory and inhibitory odor responses, while CS-activity was mainly unaffected. b. Percentage of CS+ and CS- responsive neurons, divided into novice and expert states. No change in the percentage of CS+ responding was observed upon OFC suppression. Compared to Extended Data Fig. 5e. c. The average PSTH of all CS+ and CS- odor-responding neurons in novice and expert states recorded with OFC silencing, showing no difference in response amplitudes between the odors. d. The mean firing rates (Z-scored) for CS+ and CS- odor responding neurons for novice and expert, and excitatory and inhibitory odor responses. e. Similar to **c,** for the A and B context odors f. Similar to **d**, for the A and B context odors.

**Extended Data Fig. 9:**
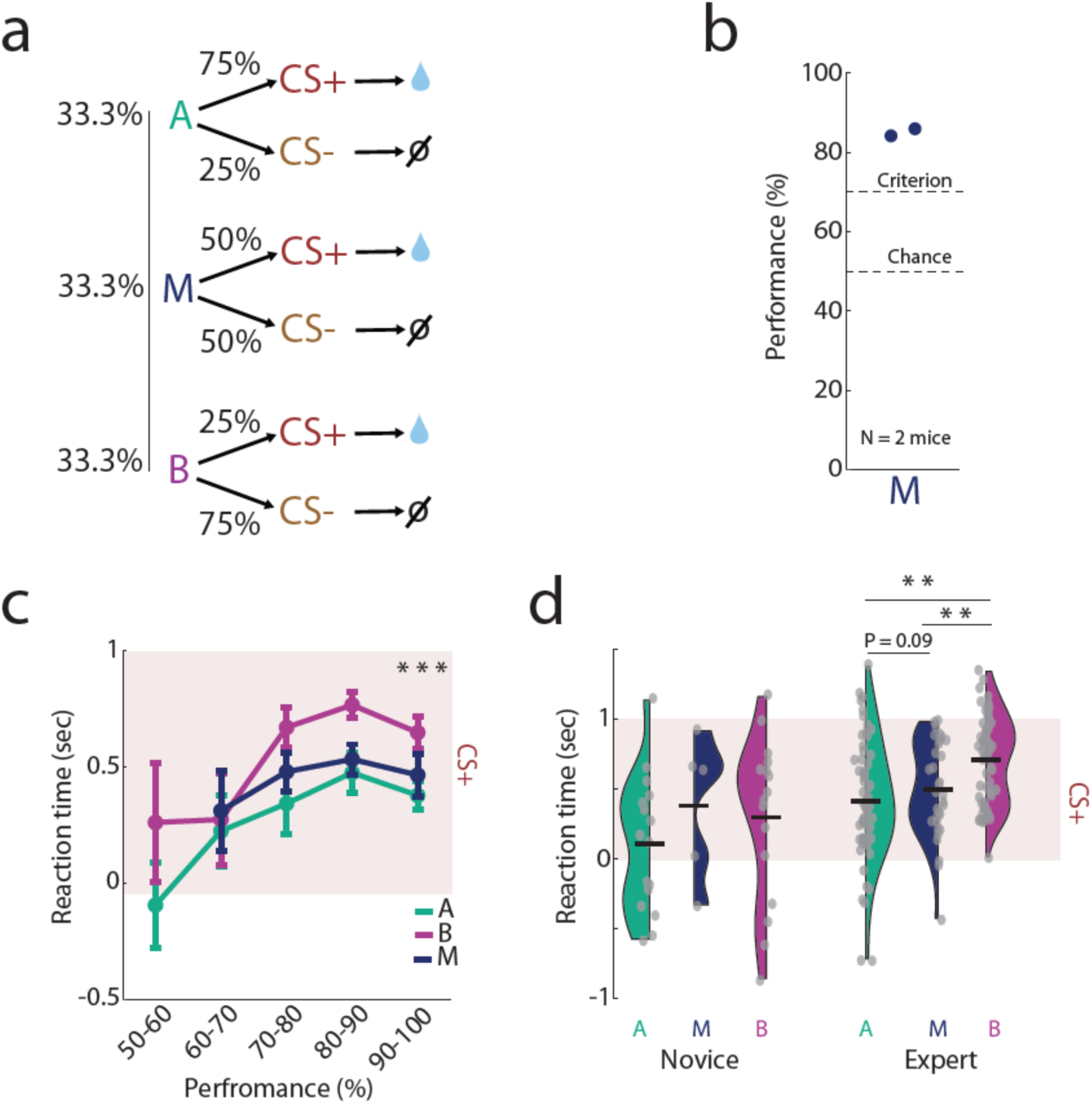
Analyses of behavioral task performance with M condition. a. Experiment contingencies. Mice learned three contexts, A, B, and M. Context M predicted the occurrence of CS+ and CS- with equal probability (N = 2 mice). b. The average discrimination success rate in the M condition in the expert state. c. Reaction times as a function of task performance. Data in the M condition (blue, N = 37 sessions) is overlaid on the data in the A and B conditions (N = 9 mice, 78 sessions, ***P < 0.001, repeated measures ANOVA over expert reaction times, performance > 70%). d. Mean reaction times in A, M, and B conditions for novice (left) and expert (right) states. Post-hoc analyses of the repeated measures ANOVA.

